# Activation of the membrane-bound Nrf1 transcription factor by USP19, a ubiquitin-specific protease C-terminally tail-anchored in the endoplasmic reticulum

**DOI:** 10.1101/2020.10.05.326363

**Authors:** Shaofan Hu, Yuancai Xiang, Lu Qiu, Meng Wang, Yiguo Zhang

**Affiliations:** The Laboratory of Cell Biochemistry and Topogenetic Regulation, College of Bioengineering and Faculty of Medical Sciences, Chongqing University, No. 174 Shazheng Street, Shapingba District, Chongqing 400044, China; Department of Biochemistry and Molecular Biology, College of Basic Medical Sciences, Southwest Medical University, Sichuan, 646000, China; School of Life Sciences, Zhengzhou University, No. 100 Kexue Avenue, Zhengzhou 450001, Henan, China

**Keywords:** Nrf1, Nrf2, USP19, proteasome, ubiquitination, deubiquitination, endoplasmic reticulum

## Abstract

The membrane-bound transcription factor Nrf1 (i.e., encoded by *Nfe2l1*) is activated by sensing glucose deprivation, cholesterol excess, proteasomal inhibition and oxidative stress, and then mediates distinct signaling responses in order to maintain cellular homeostasis. Herein, we found that Nrf1 stability and transactivity are both enhanced by USP19, a ubiquitin-specific protease tail-anchored in the endoplasmic reticulum (ER) through its C-terminal transmembrane domain. Further experiments revealed that USP19 directly interacts with Nrf1 in proximity to the ER and topologically acts as a deubiquitinating enzyme to remove ubiquitin moieties from this protein, and hence allows it to circumvent the potential proteasomal degradation. Such USP19-mediated effect takes place only after Nrf1 is retrotranslocated by p97 out of ER membranes to dislocate the cytoplasmic side. Conversely, knockout of USP19 causes significant decreases in the Nrf1 abundance and its active isoform entering the nucleus, resulting in down-regulation of its target proteasomal subunits. This led to a modest reduction of *USP19*^−/−^-derived tumor growth in xenograft mice, when compared with wild-type controls. Altogether, these demonstrate that USP19 serves as a novel mechanistic modulator of Nrf1, but not Nrf2, enabling Nrf1 to be rescued from putative ubiquitin-directed ER-associated degradation pathway. In turn, our additional experimental evidence has unraveled that transcriptional expression of endogenous *USP19* and its promoter-driven reporter genes is differentially regulated by Nrf2, as well by Nrf1, at distinct layers within a complex hierarchical regulatory network.

## Introduction

Nrf1 was identified as an endoplasmic reticulum (ER) membrane-bound transcription factor (1,2), that belongs to the Cap’n’Collar (CNC) basic-region leucine zipper (bZIP) family (3,4). This family comprises nuclear factor-erythroid 2 (NF-E2) p45 subunit and related Nrf1, Nrf2, Nrf3, Bach1 and Bach2 in vertebrates, the *Caenorhabditis elegans* protein Skn-1 and the founding *Drosophila melanogaster* Cnc, and also include another early-evolved subgroup of Nach proteins (5). They are essential for transcriptional regulation of distinct subsets of critical cognate genes responsible for homeostasis, development, health and disease (6-9). Such target genes encompass antioxidant response elements (AREs, 5’-TGAC/GnnnGC-3’) and/or other *cis*-regulatory homologues (e.g., AP1-binding site, 5’-TGAC/GTCA-3’) in their promoter regions. In mammals, Nrf1 and Nrf2 are two principal CNC-bZIP factors; either can directly bind ARE-driven genes through distinct functional heterodimers with a partner of small Maf (sMaf) or other bZIP proteins (e.g., AP1 and ATF4). Furtherly, gene-targeting experiments have revealed that Nrf1 and Nrf2 fulfill distinct biological functions through regulating different subsets of ARE-driven genes (4,10). Such distinctions between both CNC-bZIP factors are dictated by their different temporal-spatial processing mechanisms.

Accumulating evidence has also demonstrated that Nrf1 acts as an important ER sensor for intracellular redox, glucose, protein and lipid changes (11-13). In response to those biological cues, the ER-resident Nrf1 is topologically dislocated across membranes into extra-ER subcellular compartments, where this protein is subjected to its selective post-translational processing (e.g., deglycosylation, deubiquitination, and juxtamembrane proteolysis) to yield a mature CNC-bZIP factor before transactivating cognate genes (e.g. those encoding proteasomal subunits, antioxidant proteins and detoxifying enzymes). By contrast, the water-soluble Nrf2 is segregated primarily in the cytoplasm by Keap1, an adaptor subunit of the Cullin 3-based E3 ubiquitin ligase that targets this CNC-bZIP protein to ubiquitin-mediated proteasomal degradation (7). Since Keap1 also acts as a key sensor for oxidative and electrophilic stress, it allows for dissociation of Nrf2 stimulated by redox stress, so that the CNC-bZIP factor is released and then translocated into the nucleus, before transactivating target genes that are involved in antioxidant, detoxification and cytoprotective adaptation.

Such distinct tempo-spatial processing mechanisms of between Nrf1 and Nrf2 are attributable to their different structural domains. Of note, extra NTD (N-terminal domain) and NST (asparagine/serine/threonine-rich) domains are present in Nrf1, but not in Nrf2 (4). Within NTD, the NHB1 (N-terminal homology box 1) signal peptide of Nrf1 allows it to be anchored in a proper topology within and around ER membranes, whilst its NHB2 (N-terminal homology box 2)-adjoining peptide is responsive to the topobiologically-regulated juxtamembrane proteolytic processing of this CNC-bZIP protein (14,15). Once Nrf1 is anchored within the ER, its NST domain is partitioned in the lumen, in which it is glycosylated to become an inactive glycoprotein and thus protected by membranes (1). Only when it is required, some ER luminal-resident domains of the CNC protein are allowed for dynamic retro-translocation into extra-ER compartments, which is driven predominantly by p97/VCP (16). In this topovectorial process, Nrf1 is losing the protection by membranes, such that its deglycosylated protein enables to be processed to yield a mature CNC-bZIP factor or otherwise degraded by proteasomes (17).

During dynamic dislocation of Nrf1 from the ER lumen, it is postulated to be ubiquitinated by the ER-associated E3 ubiquitin ligase Hrd1 (12,14,18), which can also trigger protein retro-translocation across membranes into the cytoplasm (19,20). Thus, ubiquitin-labeled Nrf1 is recognized and degraded by proteasomes. However, the ubiquitination of Nrf1 may also be reversible, as most regulatory modifications of other proteins described by (21). Such being the case, it is inferable that ubiquitinated Nrf1 enables to be targeted for deubiquitination by an ubiquitin-specific protease (USP) and/or other deubiquitinating enzymes (DUBs), so that this CNC-bZIP protein is circumvented from the ubiquitin-mediated proteasomal degradation. Amongst the USP/DUB family, USP15 was reported to enable for deubiquitination of Keap1, so as to efficiently incorporate into the Keap1-Cul3-E3 ligase complex and thus enhance the enzymatic activity to increase Nrf2 degradation, but with a concomitant reduction in profiling Nrf2-target gene expression (22). Later, USP15 was also showed to activate Nrf1 in the nucleus by stabilizing the latter CNC-bZIP factor through their physical interaction and ensuing deubiquitination (23). Taken together, USP15 can negatively regulate Nrf2 through deubiquitination of Keap1, but also directly activates Nrf1-mediated expression of a *PSMA4*-derived ARE-luciferase reporter and endogenous proteasomal activity. In turn, Nrf1 was demonstrated to transactivate expression of *USP9x, USP14* and other *USP/DUB* genes, in addition to those encoding proteasomal subunits, p97 and their co-factors (16,24). Nonetheless, it is unknown whether the membrane-bound Nrf1 is regulated by one ER-resident USP enzyme [i.e., USP19, USP30 or USP48, that were identified by (25)].

Notably, USP19 has been showed to rescue the substrates of ER-associated degradation (ERAD) by removing their ubiquitin chain in the unfolded protein response (UPR) (25). Meanwhile USP19 also participates in the unconventional secretion to export misfolded cytosolic (e.g., neurodegenerative disease-causing) proteins into extracellular space (26). In this protein-disposing mechanism, USP19 can bind HSP70/HSC70 and also act as upstream of HSC70 and DNAJC5 (a membrane-associated chaperone in late endosomes and lysosomes). Thus, USP19 is involved in regulation of cell cycle progression, DNA damage repair, apoptosis and autophagy (27,28). Here, we found that USP19 interacts with Nrf1 to enhance this CNC-bZIP stability and its transcriptional activity. This is likely due to the experimental observation that a half-life of Nrf1 is significantly prolonged by over-expression of USP19, but shortened in *USP19*^−/−^ cells. Such effects of USP19 on Nrf1 are nearly completely prevented by mutation of all six putative ubiquitination sites within NTD and AD1 (acidic domain 1) of this CNC-bZIP protein. Loss of USP19 can also cause a reduction in Nrf1 abundances to enter the nucleus, along with reduced expression of its target proteasomal subunits, whereas Nrf2 was almost unaffected. Their ensemble effects are manifested by a modest reduction of *USP19*^−/−^-derived tumor growth in xenograft model mice. Overall, these findings demonstrate that USP19 serves as a novel modulator of Nrf1, but not Nrf2, to mediate its target gene expression.

## Results

### USP19 enhances the protein abundance of Nrf1 and its transcriptional activity

The DUB/USP family members are distributed throughout distinct cellular locations to exert different functions (29-31). Rather, it is worth mentioning that a subtle transmembrane (TM) structure exists in USP19, USP30 and USP48, but not USP14 or USP15, which enables each of those TM-containing proteases to be anchored within ER membranes (25). Here, the five expression constructs for USP14, USP15, USP19, USP30 and USP48 were created (Fig. 1A), to examine potential effects of these deubiquitinating enzymes on Nrf1. As shown in Fig. 1B, co-transfection of each indicated USP with Nrf1 revealed no obvious changes in the full-length Nrf1 glycoprotein, but only an extra conspicuous isoform of this CNC-bZIP protein emerged from co-expression with USP19 alone (as marked by red star), rather than from other examined USPs (Fig. 1B) or the empty vector (Fig. S1A). Such this putative deubiquitinated isoform of Nrf1 had a slightly less mass than the intact glycoprotein-A, but its molecular weight is much more than the deglycoprotein-B.

**Figure 1.**
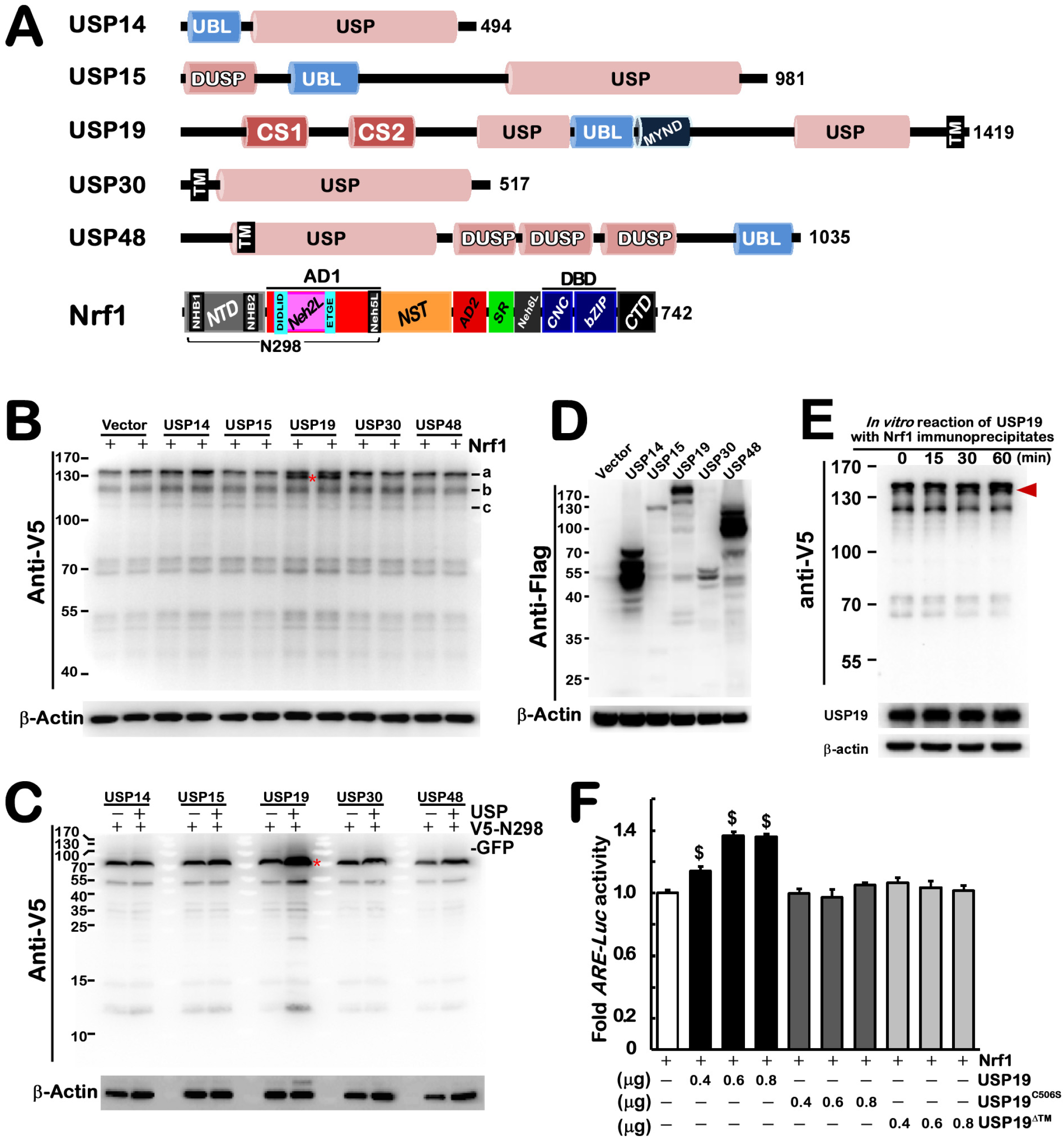
The deubiquitinating effect of USP19 on Nrf1, but not Nrf2. (A) Schematic of five ubiquitin-specific proteases (USPs) and Nrf1 with their distinct domains. Deubiquitinating enzymatic activity of the five ubiquitin-specific proteases is dictated by their founding USP domain. Besides, specific DUSP (domain present in the USP family) regulates the stability of or interact with HIF-1α, VHL or COP9-signalosome-associated cullin E3 ligase complexes, while the CS (CHORD and SGT1) domain has a feature of cochaperone interacting with multiple protein complexes. Additional locations of ubiquitin-like (UBL), zinc-finger Myeloid-Nervy-DEAF1 (MYND) and transmembrane (TM) domains are also indicated. Lastly, NTD (N-terminal domain), NHB1 (N-terminal homology box 1), NHB2 (N-terminal homology box 2), AD1(acidic domain 1), NST (asparagine/serine/threonine domain), AD2 (acidic domain 2), SR (serine-repeat domain), Neh2L (Nrf2–ECH homology 2-like), Neh6L (Nrf2–ECH homology 6-like), CNC (cap‘n’collar), bZIP (basic region leucine zipper domain) and CTD (C-terminal domain) are all indicated in Nrf1. (B, C) Expression constructs for Nrf1-V5 (B) or V5-N298-eGFP (C) plus one of deubiquitinating enzymes USP14, USP15, USP19, USP30 and USP48, were co-transfected into COS-1 cells, followed by Western blotting of the cell lysates with V5 antibody. The small letters ‘a, b, c’ represent Nrf1 glycoprotein, deglycoprotein and proteolytic proteins, respectively, [which was described in detail elsewhere (15, 16)]. Besides, a deubiquitinating protein of Nrf1 is marked by a red star. In addition, it is worth mentioning that no deubiquitinating effect of USP19 on Nrf2 was shown in supplemental Fig. S1B. (D) The expression of each Flagged USP in the above COS-1 cell lysates were examined by Western blotting with Flag antibody. (E) Nrf1-expressing lysates from COS-1 cells were allowed *in vitro* reactions with USP19 immunoprecipitates at 37°C for different times, followed by Western blotting analysis of their electrophoretic mobility of these proteins recognized by antibodies against V5 or USP19 epitopes. The red arrow indicates deubiquitinating protein of Nrf1. (F) A dose-dependent effect of USP19 on Nrf1-mediated 6×*ARE-Luc* reporter activity in COS-1 cells, that had been co-transfected for 8 h with an expression construct for Nrf1, together with different amounts (μg of cDNAs) of USP19, its mutants USP19^C506S^ or USP19^ΔTM^, plus 6×*ARE-Luc* or pRL-TK plasmids and then allowed for 24-h recovery from transfection. The resulting data were calculated as fold changes (mean ± SEM, n = 3 × 3) with significant increases ($, *p* <0.01), relative to the control value obtained from without USP19.

Further examination unraveled that the intact full-length chimeric protein V5-N298-GFP [in which N298 (i.e. the first 298 aa of Nrf1) was sandwiched between the N-terminal V5 tag and C-terminal GFP (15) as illustrated in Fig 1A] of 80-kDa, as well as its several N-terminally-cleaved polypeptides of between 55-kDa and 12.5-kDa, were markedly increased by its co-expression with USP19 only (Fig. 1C). However, no similar effects were obtained from all other four proteases examined (Figs. 1, C & D) or the empty vector (Fig. S1A). These imply that the proteolytic processing of N298 by USP19 occurs within the NTD and AD1 regions of Nrf1. Rather, such effects of USP19 on Nrf1 appeared to be not exerted on Nrf2 (Fig. S1B). This distinction may be attributable to a few of structural domains (e.g., NTD) in Nrf1, but not in Nrf2, albeit both factors shared several conserved functional domains (4).

The expression of deubiquitinated Nrf1 isoform could also be enhanced by USP19 in the *in vitro* reactions with its immunoprecipitates by Flag antibody (Fig. 1E). Moreover, the transactivation activity of Nrf1 to mediate transcriptional expression of a 6×ARE-driven luciferase reporter were incremented by USP19 in a dose-dependent manner (Fig. 1F). But, such USP19-increased transactivity of Nrf1 was completely abolished by USP19^C506S^ (a mutant of the enzymatic thiol-active site of USP19 at Cys^506^ into serine) or USP19^ΔTM^ (lacking the C-terminal TM region of this protease and hence losing its capability to be localized and anchored in the ER of cells, as determined by subcellular fractionation in Fig S1C). Overall, these indicate that USP19 can also regulate transactivation activity of Nrf1 by removing its ubiquitin-modified moieties.

### USP19 is required for the stability of Nrf1 with a prolonged half-life

To determine the effects of USP19 on Nrf1 stability and processing, the pulse-chase experiments were carried out in distinct cell lines that had been treated with 50 μg/ml of cycloheximide (CHX, to inhibit biosynthesis of nascent proteins). As anticipated, Western blotting of cell lysates showed that abundances of Nrf1α and derivative isoforms were strikingly enhanced by USP19 (Figs. 2A and S2A & B). This led to an extended course of CHX-chased immunoblots before their disappearance by 4-h treatment, when compared to control experiments without this deubiquitinating protease. The difference in the conversion between Nrf1α-derived isoforms was analyzed stoichiometrically (Fig. 2B). The stability of Nrf1α-derived glycoprotein (G), deglycoprotein (D) and processed proteins (P) (which were separated by distinct electrophoretic gels in Fig. S2A,B) was estimated by their half-lives, which were determined to be 0.40 (*vs* 0.28), 2.99 (*vs* 1.39), and 3.48 (*vs* 0.46) h after CHX treatment of COS-1 cells that had been allowed for co-expression of USP19 (*vs* not), respectively (Fig. 2B). Conversely, knockout of USP19 from HepG2 cells (that were identified in Fig. S3) resulted in obvious decreases in basal abundances of Nrf1α-derived proteins (Figs. 2C and S2C, D), along with their shortened half-lives in *USP19*^−/−^ cells (as shown graphically in Fig. 2D), by comparison with their wild-type controls.

**Figure 2.**
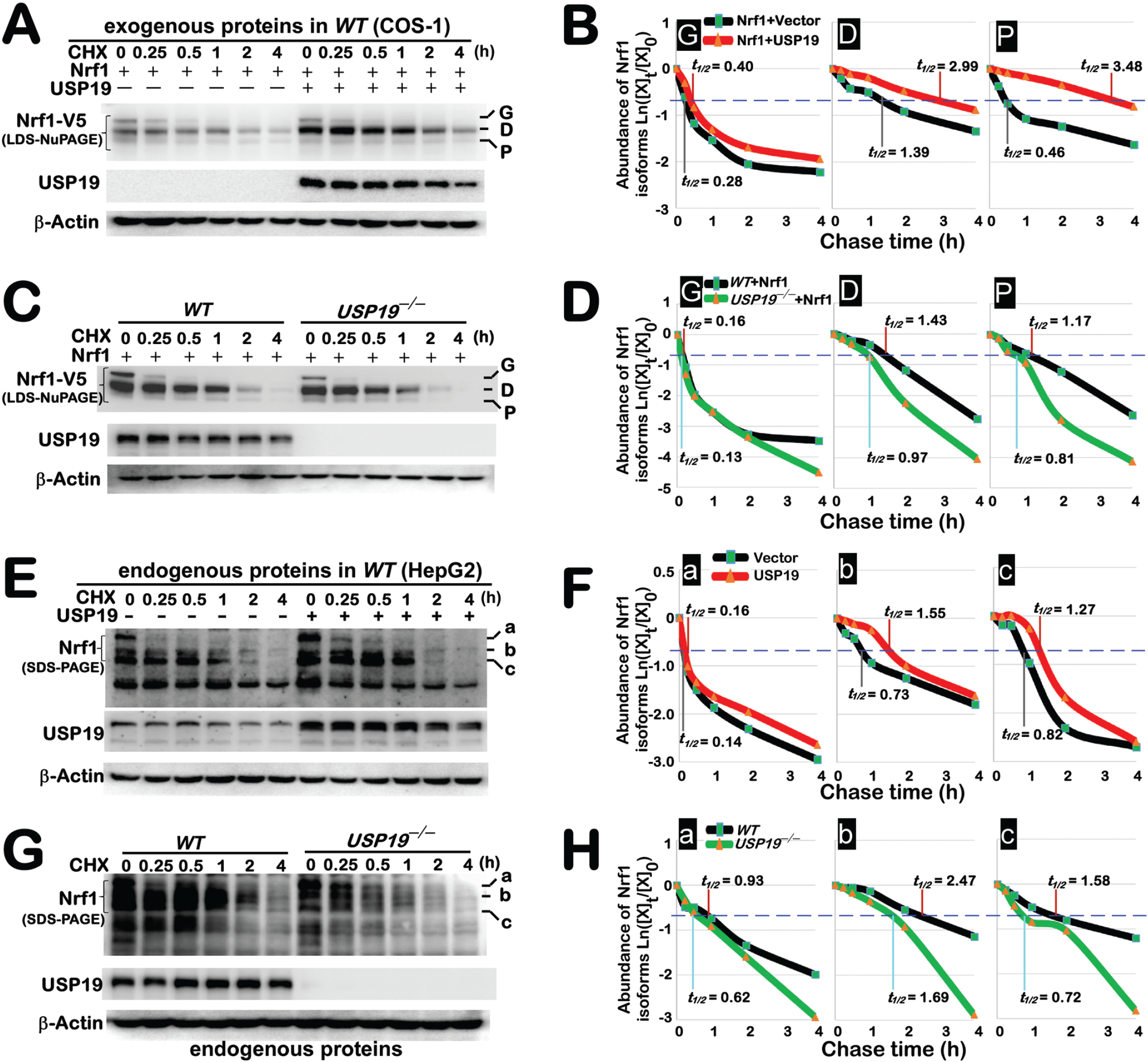
USP19 has an effect on the stability of Nrf1 isoforms with distinct half-lives. (A, B) COS1 cells expressing Nrf1 alone or plus USP19 were treated with 50 μg/ml of CHX for indicated lengths of time and then subjected to the pulse-chase analysis. The cell lysates were separated by 4-12% LDS-NuPAGE gels in pH 7.7 running buffer, and visualized by Western blotting with antibodies against USP19 or V5 tag. Subsequently, the intensity of immunoblots representing Nrf1 isoforms was quantified by Quantity One, as shown graphically (B). Of note, the inactive 120-kDa Nrf1 glycoprotein (G), its 95-kDa deglycosylated protein (D) and N-terminally-cleaved 85-kDa proteolytic protein (P) were separated by 4-12% LDS-NuPAGE gels; they were thus represented by the capital letters ‘G, D, P’, respectively. (C,D) Wild-type HepG2 and *USP19*^−/−^ cell lines were allowed for ectopic expression of Nrf1, before being subjected to the pulse-chase experiments (C) and subsequent stoichiometrical analysis of Nrf1 isoforms (D). (E, F) HepG2 cells expressing USP19 or not were subjected to CHX pulse-chase analysis of endogenous Nrf1 isoforms. The lyastes were separated by 8% SDS-PAGE gels in pH 8.3 running buffer, and visualized by immunoblotting with antibodies against Nrf1 or USP19 (E). The stoichiometrical results of Nrf1 isoforms are shown graphically (F). In addition, endogenous Nrf1 glycoprotein, deglycoprotein and proteolytic proteins were also denoted by the letters ‘a, b, c’, respectively, of which the detailed descriptions were referenced elsewhere (15, 16). (G,H) CHX-treated HepG2 (*WT*) and *USP19*^−/−^ cell lines were employed for time-course analysis of endogenous Nrf1 isoforms (G), with distinct stability as shown graphically (H).

Further examination of USP19’s effects on endogenous Nrf1 stability revealed that distinct half-lives of its three major isoforms-A, B and C were slightly prolonged by over-expression of this protease (Figs. 2E and S2E), as shown graphically (Fig. 2F). However, half-lives of these endogenous Nrf1α-derived isoforms-A, B and C were substantially shortened by knockout of *USP19*^−/−^ (Fig. 2G and S2F), which were determined to be 0.62, 1.69, 0.72 h, respectively, (Fig. 2H), when compared with their equivalent controls of wild-type cells that were estimated by half-lives of 0.93, 2.47, and 1.58 h after CHX treatment. Together, these demonstrate a requirement of USP19 for Nrf1 stability, albeit a nuance in between endogenous and ectopic USP19’s effects on Nrf1α-derived proteins exists.

### Deubiquitination of Nrf1 by its interactor USP19 in close proximity to the ER

Distinct immunofluorescent images of COS-1 cells that had been co-transfected with two expression constructs for USP19 plus Nrf1 or its mutant Nrf1^Δ2-36^ were visualized by confocal microscopy (Fig. 3A). The red fluorescent signals of USP19 are dominantly in the cytoplasm, and also superimposed with those green signals of ER-resident Nrf1. However, another green fraction of this CNC-bZIP signals were distributed in the nucleus. By sharp contrast, Nrf1^Δ2-36^ only gave predominant signals in the nucleus, where it cannot have fully engaged with the cytoplasmic USP19 signals (Fig. 3A).

**Figure 3.**
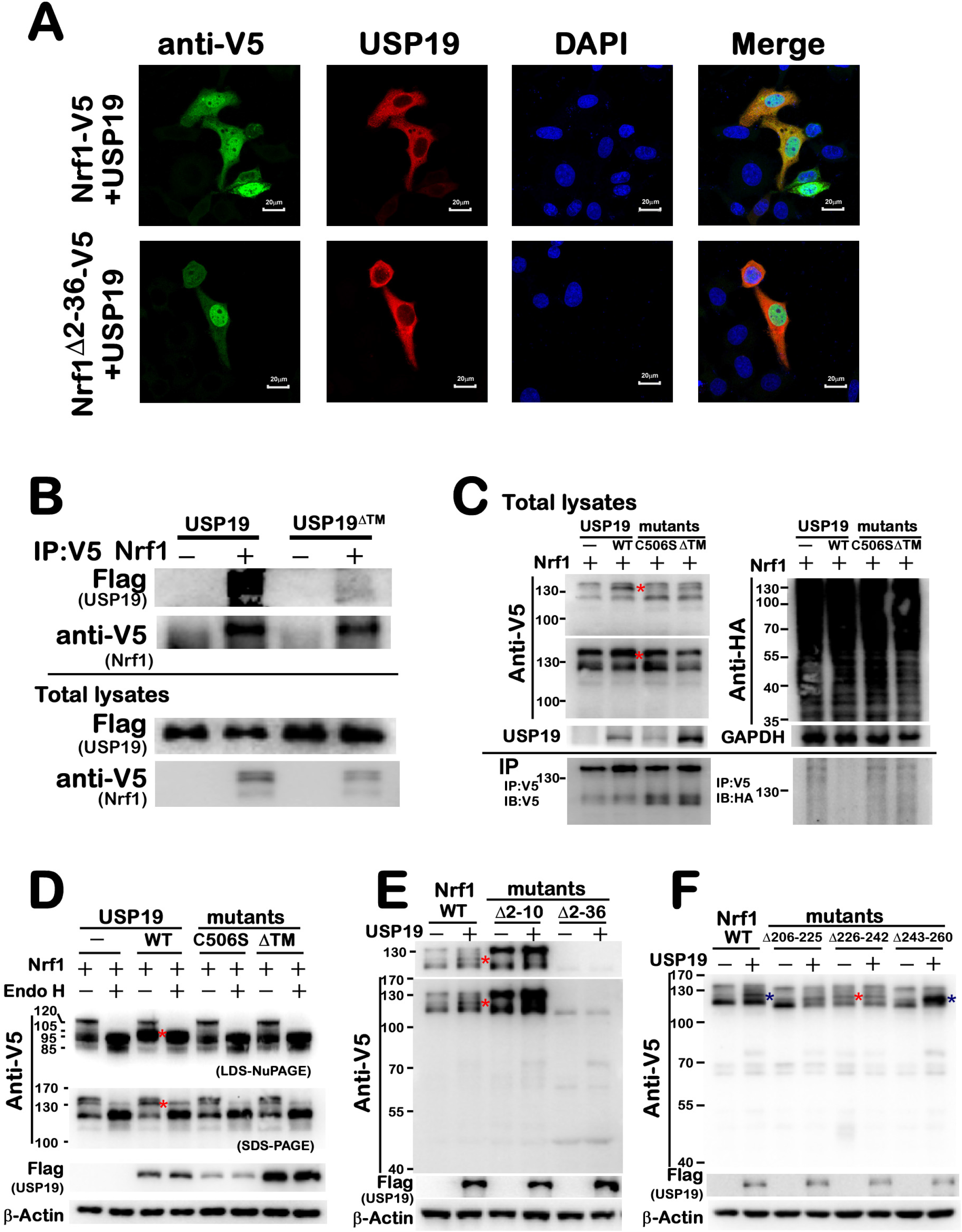
Deubiquitination of Nrf1 by its interactor USP19 in their co-location close to the ER. (A) After immunocytochemical staining of COS-1 cells co-expressing USP19 plus either Nrf1 or Nrf1^Δ2-36^ with antibodies against USP19 or V5 tag. Distinct fluorescence images were achieved and also merged (scale bar = 20 μm). The relevant subcellular fractionation of COS-1 cells co-expressing Nrf1 together with USP19 or its mutant USP19^ΔTM^ was also shown (in supplemental Fig. S1C). (B) Co-Immunoprecipitates (Co-IP) of COS-1 cells co-expressing Nrf1-V5 plus USP19-flag or its mutant USP19^ΔTM^-flag with anti-V5 antibodies were analyzed by immunoblotting with anti-Flag or anti-V5 antibodies, respectively (*upper two panels*). Besides, the inputs of whole-cell lysates were examined in parallel experiments (*lower two panels*). (C) Two distinct anti-V5 immunoprecipitates of COS-1 cells co-expressing of Nrf1-V5 and HA-Ub, plus USP19, its mutants USP19^C506S^ or USP19^ΔTM^, were visualized by immunoblotting with anti-V5 or anti-HA antibodies (*lower two panels*). The whole-cell lysates were also, together, examined (*upper two panels*). In addition, it is worth mentioning that similar CO-PI experimental analysis of endogenous proteins was also conducted (in supplemental Fig. S4,B to E). (D) COS-1 cells had been transfected with an expression construct for Nrf1, plus USP19-Flag or its mutants USP19^C506S^ or USP19^ΔTM^, and then allowed for 24-h recovery from transfection, before the cells were harvested in denatured lysis buffer. Subsequently, the cell lysates were allowed for deglycosylation reaction with Endoglycosidase H (Endo H) or not, before being separated by 8% SDS-PAGE gels or 4-12% LDS-NuPAGE gels in distinct running buffers, and then visualized by immunoblotting with antibodies against V5 tag or USP19, respectively. (E, F) Expression constructs for Nrf1 and its mutants examined were allowed for co-expression with USP19 or not with this protease, before immunoblotting with V5 or USP19 antibodies.

Co-immunoprecipitation (Co-IP) of the above-described cell lysates by V5 antibody revealed a direct interaction of Nrf1-V5 with the flagged USP19 (Figs. 3B, S4A), but not with another flagged mutant USP^ΔTM^ (Fig. 3B). The endogenous Nrf1 was also shown to directly co-immunoprecipitate with USP19 put down by an Nrf1 antibody (Fig. S4B). Further Co-IP assays of the other cell lysates, which had been co-transfected with expression constructs for Nrf1-V5 and HA-ubiquitin (Ub) together with USP19, its mutants USP19^C506S^ or USP19^ΔTM^, revealed that the abundance of putative Ub-conjugated Nrf1-V5 proteins was substantially reduced by this deubiquitinating protease USP19, rather than by its two mutants (Figs. 3C and S4C). The endogenous immunoprecipites put down by Nrf1 antibody were further visualized by anti-Ub immunoblotting, demonstrating that the reduction of ubiquitinated Nrf1 by USP19 was greatly recovered and reversed from this protease knockout in *USP19*^−/−^ cells (Fig. S4D, *lower panels*). In addition to the deubiquitinated Nrf1, direct interaction of USP19 with putative ubiquitinated Nrf1 proteins was also verified by another Co-IP assays for anti-Flag immunoprecipitates of cell lysates co-expressing Nrf1-V5 and HA-Ub along with USP19, but not with USP19^ΔTM^ (Fig. S4E, *lower panels*). Taken together, these indicate a major interaction of USP19 with low-ubiquitinated (or partial deubiquitinated) Nrf1; this is accompanied by a minor interaction of USP19 with the high-ubiquitinated Nrf1 (primarily in the USP19-deficient cell lines). Thereby, it is inferable that upon interaction of USP19 with ubiquitinated Nrf1, this may allow for rapid deubiquitination of this CNC-bZIP protein by this protease, so that Nrf1 is rescued by USP19 from putative Ub-mediated ER-associated degradation pathway and its resulting protein stability is hence enhanced.

Further experiments confirmed that USP19-deubiquitinated Nrf1 isoform (marked by red star) was abolished by its mutants USP19^C506S^ or USP19^ΔTM^ (Fig. S4F). In the meantime, *in vitro* deglycosylation assays unraveled that such a deubiquitinated isoform of Nrf1 was remarkably enhanced after being treated by USP19, but not by USP19^C506S^ or USP19^ΔTM^ and its molecular weight was obviously located between both glycoprotein and deglycoprotein of Nrf1 (Figs. 3D and S4G,H). Besides, USP19 also enhanced abundances of the intact 80-kDa V5-N298-GFP and its N-terminally-cleaved 55-kDa polypeptide, but this enhancement was also reduced by USP19^ΔTM^ (Fig. S5A). These demonstrate that deubiquitination of Nrf1 (within its N298 portion) by USP19 occurs in the closer proximity to their co-tethered ER membranes; this is also supported by the subcellular fractionation of USP19^ΔTM^ lacking its ability to be anchored in the ER (Fig. 1C). Collectively, these implicate that only membrane-tethered USP19 exerts its enzymatic activity to mediate deubiquitination of Nrf1 (and its N298 fusion proteins) localized within and around the ER or close proximity to the organelle membrane. This notion is further evidenced by loss of its ER-targeting signal to yield the mutant Nrf1^Δ2-36^ (Fig. 3E). However, it was, to our surprise, found that such deubiquitinated Nrf1 isoform appeared to be constructively abolished by Nrf1^Δ2-10^ mutant (lacking the lysine-rich n-region of its ER-targeting signal) (Fig. 3E), but also constitutively emerged from another mutant Nrf1^Δ226-242^ (lacking a putative Keap1-binding ETGE motif) (Figs. 3F & 4A). The presence or absence of deubiquitinated Nrf1 mutant isoform appeared to be largely unaffected by USP19, implying that there exists another intrinsic modification of Nrf1 by ubiquitination prior to USP19-mediated deubiquitination.

### Distinct contributions of various N298 peptides to USP19-enhanced abundances of Nrf1 and its processing

To gain in-depth insight into distinct contributions of various N298 peptides (Fig. 4A) to USP19-enhanced stability of Nrf1 and its proteolytic processing, here we created a series of expression constructs by internal mutagenesis mapping of the N298 region within V5-N298-GFP (as referenced (15)). As anticipated, abundances of intact full-length V5-N298-GFP of 80-kDa and its three major N-terminally-cleaved polypeptides of 55-kDa, 35-kDa and 12.5-kDa were strikingly enhanced by USP19 (Figs. 4B, 4C and S5B). However, such USP19-triggered effects were abolished by two mutants V5-N298^Δ2-10^-GFP (lacking aa 2-10 of Nrf1, that cover the n-region its ER-targeting signal facing the cytoplasmic side, Fig. 4B) and V5-N298^Δ226-242^-GFP (lacking aa 226-242 of Nrf1 containing the ETGE motif in AD1, Fig. 4C), but almost unaffected by other mutants examined (Figs. 4B, 4C and S5B). These imply that both regions of aa 2-10 and 226-242 within Nrf1 may serve as USP19-binding sites or its enzyme-targeting sites. This notion is also evidenced by the findings that lack of aa 2-10 in Nrf1^Δ2-10^ led to disappearance of deubiquitinated Nrf1 isoform (Fig. 3E), while loss of aa 226-242 in Nrf1^Δ226-242^ gave rise to a constitutive deubiquitinated Nrf1 isoform (Fig. 3F). Such disappearance of deubiquitinated Nrf1 isoform and its constitutive emergence were also unaffected by the presence and absence of USP19.

**Figure 4.**
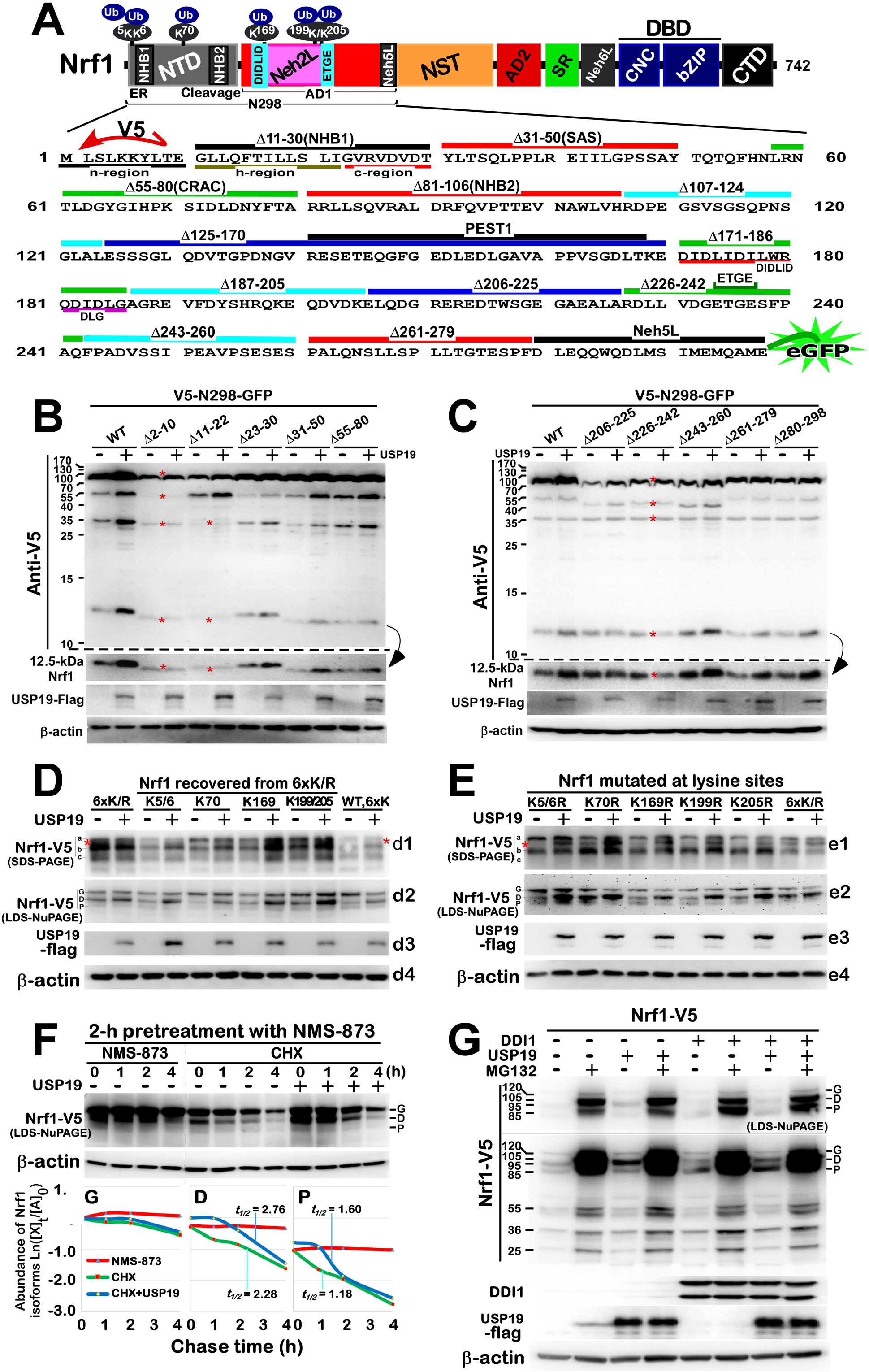
Putative deubiquitination of Nrf1 by USP19 contributes to its stability and processing. (A) Schematic of distinct functional domains in Nrf1 and its mutants of relevant regions within NTD and AD1. Within its NTD, three putative ubiquitin-conjugated lysines are situated close to its NHB1-adjoining ER-targeting or NHB2-adjacent cleavage sequences. Another three lysines are located within AD1. The N298 portion of fusion proteins is composed of both NTD and AD1 portions, in which distinct regions or motifs covering those amino acid residues as underlined were deleted from V5-N298-eGFP to yield a series of mutant fusion proteins as deciphered respectively. (B,C) COS-1 cells co-expressing of V5-N298-GFP or its deletion mutants together with USP19 or not were further evaluated by Western blotting with V5 or USP19 antibodies (also see Figure S5B). (D) Total lysates of COS-1 cells that had been transfected with expression constructs for wild-type Nrf1 (WT, 6×K), its mutant Nrf1^6×K/R^ (i.e., 6×K/R) or one of those recovered variants (i.e., K5/6, K70, K169 and K199/K205) from the Nrf1^6×K/R^ mutant, plus USP19 or not, were separated by 8% SDS-PAGE gels or 4-12% LDS-NuPAGE gels in distinct running buffers, and visualized by immunoblotting with antibodies against V5 tag or USP19, respectively. (E) Western blotting of COS-1 cell lysates expressing one of distinct lysine-to-arginine mutants of Nrf1 (i.e. K5/6R, K70R, K169R, K199R, K205R and 6×K/R) alone or plus USP19 was carried out as described above. (F) COS-1 cells expressing Nrf1 alone or plus USP19 were pretreated for 2 h with NMS-873 (10 µmol/L) and then treated with CHX (50 µg/ml) for indicated lengths of time, before being harvested. The cell lysates were resolved by 4-12% LDS-NuPAGE gels and visualized by Western blotting as described above. Subsequently, the intensity of three major immunoblots representing Nrf1 polypeptides was quantified, as shown graphically (*bottom*). (G) Western blotting analysis of COS-1 cells that had been allowed for expressiong of Nrf1 alone or plus USP19 and/or DDI1, and then treated for 2 h with MG132 (+ 5 µmol/L) or not (–) was carried out as described above.

Apart from aa 2-10 and 226-242 regions of Nrf1, its mutants of other remaining peptides within N298 were also involved in regulation of this CNC-bZIP factor by USP19 (Figs. 4B, 4C and S5B), no matter whether they are selectively processed through topovectorially-regulated juxtamembrane proteolysis (15). Of note, aa 11-22 of Nrf1 (i.e., covering the core h-region of NHB1 signal peptide that enables this CNC-bZIP protein to anchor within the ER membranes) were deleted to yield V5-N298^Δ11-22^-GFP, such that its smaller N-terminally-cleaved polypeptides of between 35-kDa and 12.5-kDa, but not its longer 55-kDa polypeptide, appeared to be unaltered by USP19 (Fig. 4B). By sharp contrast, the N-terminal 12.5-kDa polypeptide of Nrf1 was substantially abolished by deletion of its NHB2-adjoining peptides (to yield two mutants V5-N298^Δ81-106^-GFP and V5-N298^Δ107-124^-GFP), but basal abundances of their N-terminally-processed 35-kDa polypeptides were almost unaffected by USP19 (Fig. S5B). In addition, deletion of the DIDLID/DLG-adjoining peptides of aa 125-170 and 171-186 yielded two unstable mutants of V5-N298^Δ81-106^-GFP and V5-N298^Δ107-124^-GFP, such that both fusion proteins were subjected to rapidly proteolytic processing to give rise to a major 70-kDa or 40-kDa polypeptides, respectively, along with those continuously processed small polypeptides, but they were still enhanced by USP19 (Fig. 4C). Collectively, these demonstrate that such discrete effects of USP19 on basal abundance of Nrf1 and its N-terminal processing are modulated selectively through its membrane-topological mechanism, as described by (15,32,33).

### Putative deubiquitination of Nrf1 by USP19 contributes to its stability and processing

Ubiquitination of Nrf1 was considered to occur prior to, and be essential for, the proteolytic processing of this CNC-bZIP protein to yield various lengths of cleaved polypeptides (16). But, our previous work had revealed that putative ubiquitination of Nrf1 (at Lys^5,6,70^ in its NTD and/or Lys^169,199,205^ in its AD1, Fig. 4A) is not a prerequisite for retro-translocation of this CNC-bZIP protein by p97, before being subjected to its selective processing by cytosolic proteases (15). These controversial results have led us to further test what contribution of Nrf1 deubiquitination to its proteolytic processing is made *de facto*. As anticipated, co-expression of both USP19 and Nrf1 enabled this CNC-bZIP protein to give rise to an obvious deubiquitinated isoform (resolved by 10% SDS-PAGE gels in pH8.3 Tris-Glycine running buffer, Figs. 1B & 4D1). Such a similar USP19-deubiquitinating isoform of Nrf1 had also emerged directly from the mutant Nrf1^6×K/R^ (in which all six potential ubiquitin-conjugated lysines were substituted by arginines), and its abundance were unaffected by this protease (Figs. 4D1, 4E1). However, the deubiquitinating isoform of Nrf1^6×K/R^ was, to varying degrees, diminished or even abolished by recovery of the indicated lysines from this mutant (e.g., Nrf1^K70+5×K/R^ made from the K70 recovery from Nrf1^6×K/R^) (Figs. 4D1 & S6A). Further, Nrf1^K70R^, but not Nrf1^K5/6R^, Nrf1^K169R^, Nrf1^K199R^ or Nrf1^K205R^, gave rise to a similar deubiquitinated isoform to that arising from Nrf1^6×K/R^, but also its abundance was substantially highlighted by USP19 (Figs. 4E1 & S6C). By contrast, the putative USP19-deubiquitinating isoform of Nrf1^K5/6R^, Nrf1^K169R^, Nrf1^K199R^ or Nrf1^K205R^ was presented only after co-expressing this protease. Collectively, these indicated that deubiquitination of Nrf1 by USP19 occurs at all six lysine residues, of which the Lys^70^ residue serves as a key site for this protease.

Further examination of protein separation by 4-12% LDS-NuPAGE in pH7.7 MOPS running buffer) revealed that USP19 enables wild-type Nrf1 glycoprotein to be reduced, and instead of its deglycoprotein being enhanced by this protease (Figs. 4D2 and S6B). By contrast, the glycoprotein abundances of Nrf1^6×K/R^ and other mutants appeared to be not or less affected by USP19, but their deglycoprotein abundances were increased to varying extents (Figs. 4D2, 4E2, S6B and S6D). Of note, a relative stronger deglycoprotein of Nrf1^K70R^ was generated in the absence of USP19, but a similar deglycoprotein of Nrf1^K70+5×K/R^ was almost prevented by the K70 recovery from Nrf1^6×K/R^, even though their abundances were still augmented by USP19 (Figs. 4D2 & 4E2). Together, these suggest that the stability of Nrf1 and its processing (particularly when modified at K70) may be altered by USP19-mediated deubiquitination.

To clarify a role of USP19 in p97-driven retrotranslocation of Nrf1 and its subsequent conversion by proteolytic processing by cytosolic proteases, we performed a pulse-chase experiment of COS-1 cells that had been allowed for expression of Nrf1 or co-expression with USP19, and then pretreated with the p97-specific inhibitor NMS-873 for 2 h, before addition of 50 μg/ml of CHX. As excepted, inhibition of p97-mediated repositioning of Nrf1 by NMS-873 into extra-ER compartments led to a marked accumulation of this CNC-bZIP glycoprotein, as accompanied by a gradual reduction of its deglycoprotein (Figs. 4F & S6E, *lanes 1*–*4*). Subsequently, the recovery of p97 from its inhibition by NMS-873, at the same time when the nascent protein synthesis were blocked by CHX, rendered the existing ER-located Nrf1 to be dynamically dislocated into the cytoplasmic side of membranes, whereupon it was then subjected to its successive processing. The results revealed that time-lapse conversion of Nrf1 glycoprotein was unaffected by USP19 (Figs. 4F & S6E, *lanes 5*–*12*), as showed graphically (Fig. 4F, *lower left panel*). By striking contrast, turnover of Nrf1 deglycoprotein and its major processed isoform was prolonged by USP19, which was determined by obvious changes in their half-lives that were respectively extended from 2.28 to 2.76 and from 1.18 to 1.60 h after CHX treatment, when compared to those equivalents without USP19 (Fig. 4F, *lower middle and right panels*). This demonstrates that deubiquitination of Nrf1 by USP19 occurs only after dislocation of this CNC-bZIP protein to yield its deglycoprotein and rescues from being degraded by proteasomes and/or other cytoplasmic proteases (e.g. DDI1).

Co-transfection experiments of Nrf1 with USP19 or DDI1 alone or both together unraveled that this CNC-bZIP protein is subjected to the putative proteolytic processing by DDI1 (Fig. 4G). Notably, USP19 enhanced abundances of Nrf1 deglycoprotein and processed protein. However, this processed protein was increased by DDI1, as accompanied by an instead decrease in deglycoprotein of Nrf1 (Fig. 4G).

### USP19 exerts a biological function required for Nrf1-governed proteostasis and tumorigenesis

Since deubiquitination of Nrf1 by USP19 confers it to be rescued from the estabolished ubiquitin proteasomal degradation, this CNC-bZIP factor is thus accumulated and also allowed for transcriptional regulation of proteasomal subunits by a similar way to the ‘bounce-back’ response to limited proteasomal inhibition (12). Here, to explore such a similar biological role of USP19 in regulating Nrf1, we examined distinct transcriptional expression of its target proteasomal (*PSM*) subunits in *USP19*^−/−^, *Nrf1α*^−/−^ and wild-type (*WT*) cells. As showed in Fig. 5A, mRNA expression levels of all six examined genes (*PSMA1, PSMB5, PSMB6, PSMB7, PSMC6* and *PSMD12*) were significantly suppressed in *USP19*^−/−^ cells to varying extents, that are roughly similar to or slightly less than those measured from *Nrf1α*^−/−^ cells. Of note, endogenous Nrf1 deubiquitinated isoform was diminished in *USP19*^−/−^ cells, but recovered by restoration of USP19 (Fig. 5B). Further examination revealed largely similar decreases in basal protein expression levels of three core enzymatic subunits (*PSMB5, PSMB6, PSMB7*) were determined in *USP19*^−/−^ and *Nrf1α*^−/−^ cell lines (Fig. 5C). Conversely, forced expression of USP19 only led to modest increases in mRNA expression of some proteasomal subunits (Fig. S7A).

**Figure 5.**
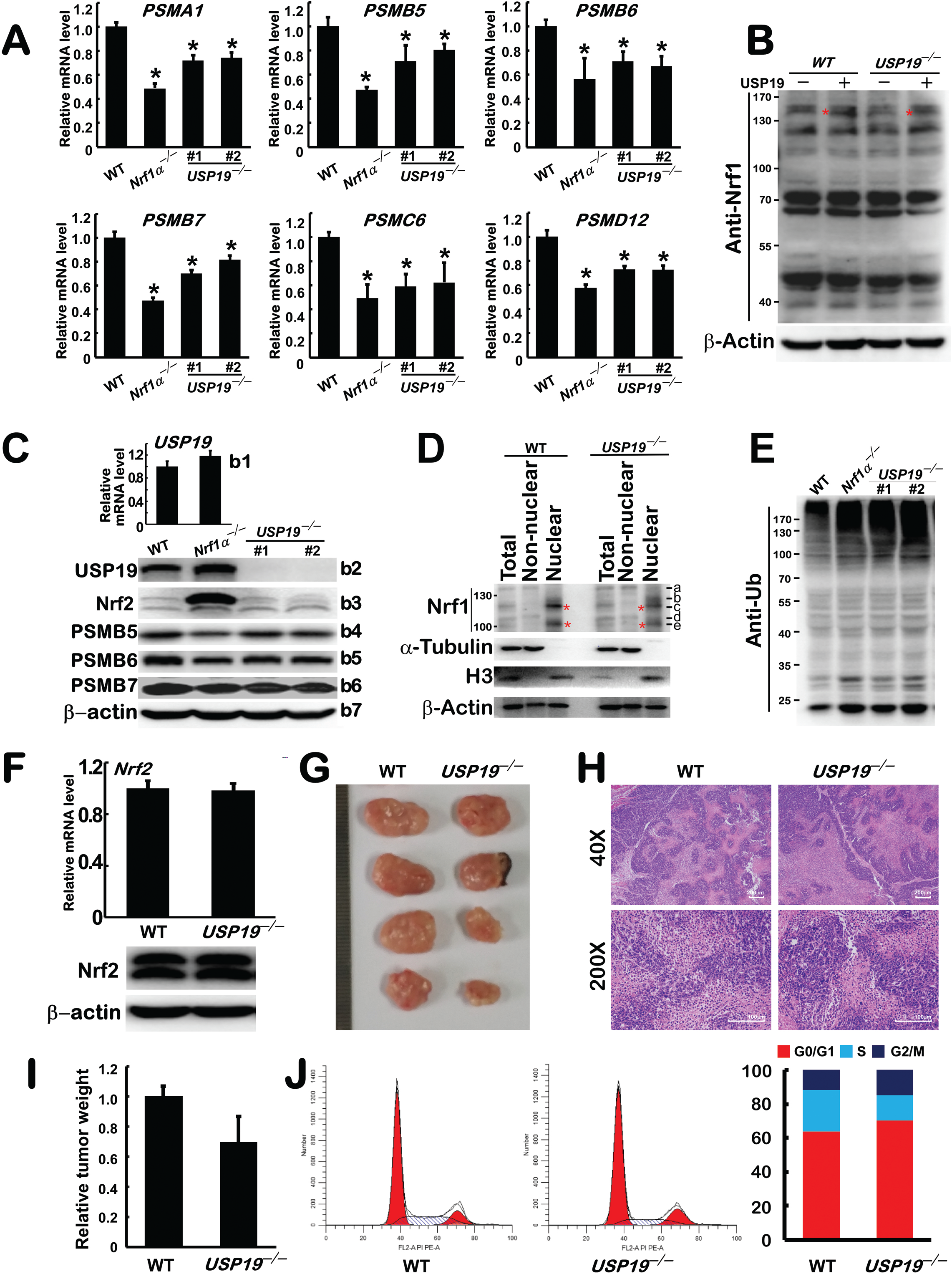
A biological function of USP19 is required for Nrf1-governed proteostasis and tumorigenesis. (A) Transcriptional expression of Nrf1-target proteasomal subunits *PSMA1, PSMB5, PSMB6, PSMB7, PSMC6* and *PSMD12* in *WT, Nrf1a*^−/−^ and *USP19*^−/−^ cell lines was examined by real-time qPCR. The resulting data are shown as fold changes (mean ± SEM, n= 3 × 3) with significant decreases (* *p* < 0.01) as compared to the controls. (B) Expression construct of USP19 or an empty vector was transfected into wild-type HepG2 and *USP19*^−/−^ cell lines, followed by Western blotting of their cell lysates with Nrf1 antibody. (C) *Top panel* shows real-time qPCR analysis of mRNA expression levels of *USP19* in *WT and Nrf1 a* ^−/−^ cells. *The lower panels* manifest Western blotting of *WT, Nrf1a*^−/−^ and *USP19*^−/−^ cell lysates with distinct antibodies against USP19, Nrf2, PSMB5, PSMB6 or PSMB7, respectively. (D) Subcellular fractions of *WT* and *USP19*^−/−^ cells were determined by Western blotting with the indicated antibodies. (E) Potential ubiquitinated proteins in *WT, Nrf1a*^−/−^ and *USP19*^−/−^ cells were analyzed by anti-UB immunoblotting. (F) Both mRNA and protein expression levels of Nrf2 in *WT* and *USP19*^−/−^ cells were detected as described above. (G) Two distinct phenotypes of xenograft tumors in nude mice were derived from subcutaneous inoculation *WT* and *USP19*^−/−^hepatoma cell lines in nude mice. (H) The pathohistological photographs of the above xenograft tumors were achieved after HE staining. (Scale bar = 200 *µ*m in ×40 or 100 *µ*m in ×200). (I) Relative weights of xenograft tumors derived from *WT* and *USP19*^−/−^ cells were calculated as shown graphically. (J) Flow cytometry analysis of *WT* and *USP19*^−/−^ cell cycles was illustrated (*left panel*). The data of three independent experiments (*n* = 3) are calculated and shown as distinct columns (*right panel*).

Subcellular fractionation revealed that, apart from the full-length glycoprotein-A of Nrf1 in the cytoplasmic non-nuclear fraction, all other Nrf1-processed isoforms-B to -E was recovered predominantly in the nuclear fraction, but also obviously reduced in *USP19*^−/−^ cells, when compared with *WT* cells (Figs. 5D and S7B). Further recovery experiment showed that a decrease of endogenous Nrf1 expression in *USP19*^−/−^ cells was reversed, as accompanied by restoration of USP19 (Fig. 5B). Thereby, it is inferable that knockdown of USP19 causes a marked decrease in the processed active Nrf1 and results in reduced expression of proteasomal subunits. This is supported by further experiments, showing evident accumulation of ubiquitinated proteins to roughly similar extents in both cell lines of *USP19*^−/−^ and *Nrf1α*^−/−^ (Fig. 5E). These data indicate that the cellular proteostasis is significantly disrupted in *USP19*^−/−^ cells by dysfunction of the proteasomal proteolytic degradation of ubiquitinated proteins. Such being the case, *USP19*^−/−^ cells still displayed a weaken resistance to cytotoxicity of the proteasomal inhibitor bortezomib (BTZ), whilst *Nrf1α*^−/−^ cells were embodied with more vulnerability to BTZ, when compared with that of *WT* cells (Fig S7C).

By contrast with Nrf1, both mRNA and protein levels of Nrf2 appeared to be unaltered in *USP19*^−/−^ cells (Fig. 5F). Yet, further investigation of *USP19*^−/−^-derived tumor growth in xenograft model mice showed a modest reduction in the subcutaneous tumorigenesis by knockout of this protease, when compared to *WT* control mice (Fig. 5G). The tumor pathohistochemistry revealed a lower tumorigenicity of *USP19*^−/−^ cells than that of its parent *WT* HepG2 cells (Fig. 5H, 5I). Subsequently, flow cytometry unraveled that *USP19*^−/−^ cell cycle seemed to be arrested at its prolonged G0/G1 phase, instead of its shorter S phase (Figs. 5J), but no changes in apoptosis of this cell lines were observed by comparison with *WT* cell controls (Fig. S8A). Taken together, these results demonstrate that USP19 plays a biological role required for Nrf1-governed proteostasis and tumorigenesis.

### Transcriptional expression of USP19 is mediated by Nrf2, as well by Nrf1, at distinct layers

Interestingly, we also found a significant diminishment in mRNA and protein expression levels of USP19 in *Nrf2*^−/−^ cells (Fig. 6A). In turn, over-expression of Nrf2 enabled USP19 expression to be significantly increased (Fig 6B). This implies the transcriptional regulation of USP19 by Nrf2. To address this, a *USP19-Luc* (i.e., its gene promoter-driven luciferase) reporter was constructed and subjected to co-transfection with an Nrf2 expression plasmid. As anticipated, the results revealed that transcription of *USP19-Luc* reporter was elevated by Nrf2. Further analysis of the *USP19* gene promoter showed five ARE sites near the transcriptional start site (Fig. 6C). Any single mutation of ARE sites caused a reduction in the transcriptional activity of *USP19-Luc* reporter mediated by Nrf2 (Fig. 6C, *lower left panel*). When all five ARE sites were mutated to yield a Mut5 reporter, Nrf2 could hardly induce the activity of this mutant *USP19-Luc* reporter. By contrast, ectopic expression of Nrf1 also stimulated induction of the *USP19-Luc* gene transcription (Fig. 6C, *lower right panel*), but this induction was only partially reduced by ARE3 mutant rather than others. However, loss of *Nrf1α* also caused a modest increase in the expression of *USP19 per se* (Fig. 5C); this is hence deduced to result from aberrant hyper-expression of Nrf2 in *Nrf1α*^−/−^ cells (Fig. S8B). Therefore, it is inferable that *bona fide* transcription of *USP19* regulated predominantly by Nrf2 is suppressed by loss of Nrf1*α* to a considerable lower extent in *Nrf1α*^−/−^ cells (Fig. 6D). This notion is further supported by similar results obtained from transcriptomic analysis of both *Nrf1α*^−/−^ and *Nrf2* ^−/−^ cell lines, showing almost no changes in the mRNA expression of *USP19* in *Nrf1α*^−/−^ cells, as accompanied by a significant decrease in the *USP19* expression in *Nrf2* ^−/−^ cells (Fig. S8C), although *Nrf1α*^−/−^-led proteasomal dysfunction had resulted in the apparent accumulation of Nrf2 and USP19 (Figs. 5C and S7B). Collectively, these data demonstrate that differential expression of USP19 is mediated by such two CNC-bZIP factors Nrf1 and Nrf2 at distinct layers.

**Figure 6.**
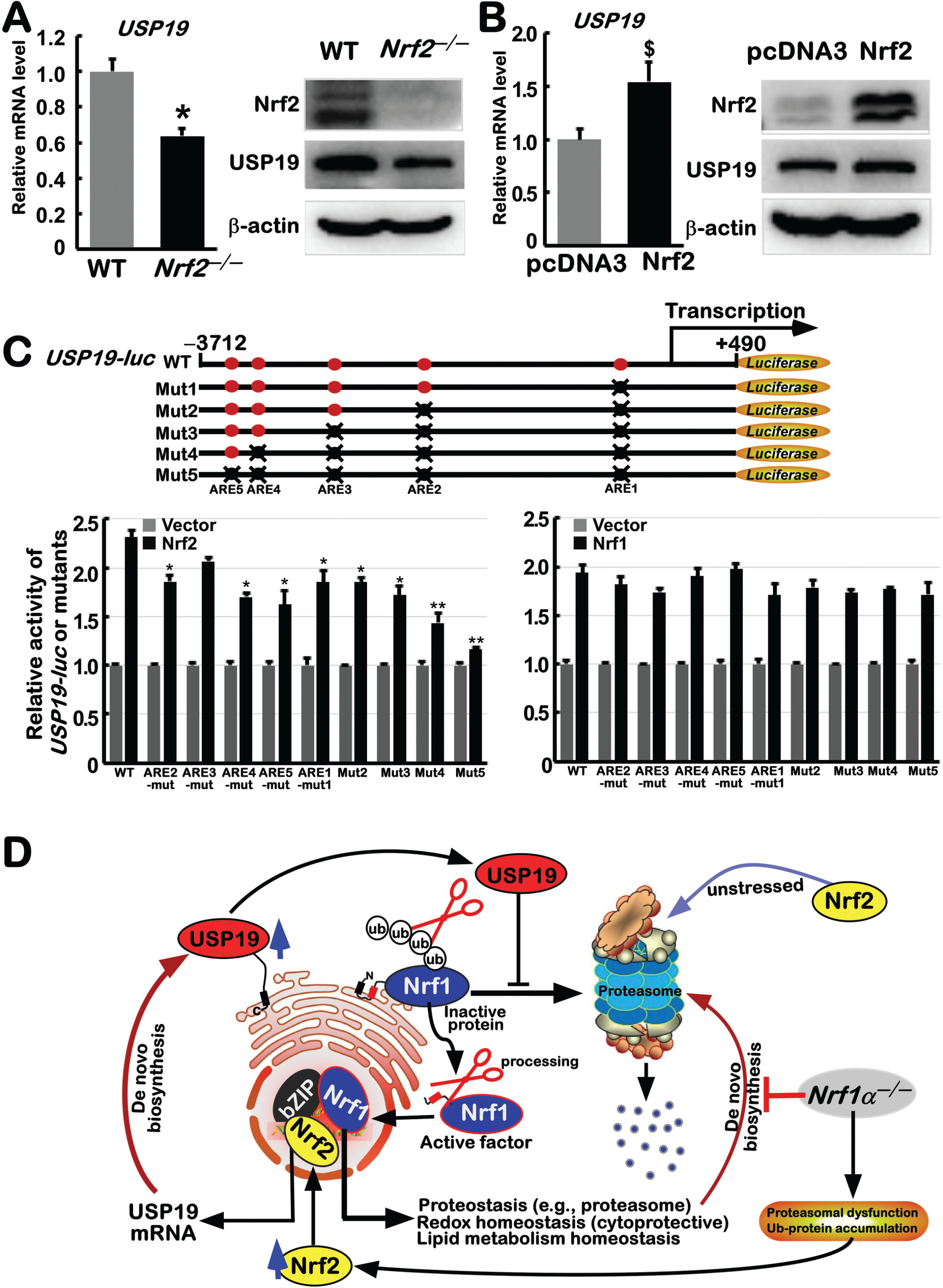
Transcriptional regulation of USP19 by Nrf2, as well by Nrf1. (A) *Top panel* shows mRNA expression levels of *USP19* in *WT* and *Nrf2*^−/−^cells, which are shown as fold changes (mean ± SEM, *n* = 3 × 3) with a significant decrease (* *p* < 0.01). Their protein expression levels were determined by Western blotting with antibodies against USP19 or Nrf2 (*middle two panels*). (B) HepG2 cells expressing Nrf2 or not were subjected to examination of *USP19* mRNA expression levels as shown as mean ± SEM (*n* = 3 × 3) with a significant increase ($, *p* < 0.01) (*upper panel*). Both protein abundances of USP19 and Nrf2 (*middle two panels*) were determined by Western blotting as described above. (C) The *upper panel* schematic shows five ARE sites in the promoter of *USP19* and also their mutants of the *USP19-luc* reporter in ARE sites. *The lower panel* shows that COS-1 cells had been co-transfected with *USP19-luc* or each of its different mutants in ARE sites, plus *pRL-TK* reporters, together with an expression construct for Nrf2 or Nrf1, or an empty pcDNA3 vector, and then allowed for 24-h recovery from transfection, before the luciferase activity was measured. The resulting data are calculated as fold changes (mean ± SEM, *n* = 3 × 3) with significant increases ($, *p* < 0.01) relative to the controls. (D) A model is proposed to provide a better explanation of the Nrf2-USP19-Nrf1-Nrf2 axes along with their inter-regulatory feedback circuit, in order to maintain the robust steady-state of cell proteostasis, beyond redox homeostasis.

Scrutiny of our transcriptomic results revealed that most of all five deubiquitinase families are regulated by Nrf1 and Nrf2 alone or both (Fig. S8C). It was found that *COPS5, PRPF8, USP2, USP39, PSMD7* and *PSMD14* were down-regulated, as accompanied by up-regulated *OTUD1, USP4, USP20, USP21* and *USP35*, in *Nrf1α*^−/−^ cells, but they were almost unaltered in *Nrf2* ^−/−^ cells, implying these 11 genes are Nrf1-targeted only. By contrast, additional 14 genes, including *UCHL5, STAMBPL, USP1, USP7, USP11, USP12, USP16, USP18, USP24, USP28, USP32, USP33, USP34* and *USP48*, were up-regulated in *Nrf2* ^−/−^ cells, but unaffected in *Nrf1α*^−/−^ cells, indicating that they are Nrf2-specific target genes. Further comparison of *USP14, USP15* and *USP25* revealed that they were up-expressed in *Nrf2* ^−/−^ cells, but down-expressed in *Nrf1α*^−/−^ cells (in which the hyperactive Nrf2 was retained), indicating that they are predominantly negatively regulated by Nrf2, rather than Nrf1. Another comparison of up-expression of *COPS6* in Nrf2-elevated *Nrf1α*^−/−^ cells, but with its down-expression in *Nrf2* ^−/−^ cells, uncovered positive regulation of this gene by Nrf2, but not Nrf1. Rather, down-regulation of *USP5* and *USP36* in both *Nrf1α*^−/−^ and *Nrf2* ^−/−^ cell lines suggested that they are transcriptionally mediated by Nrf1 and Nrf2 together. Overall, these demonstrate there exists a complex hierarchical regulatory network, which is composed of Nrf1 and Nrf2, plus USP19 and other deubiquitinases, together with the ubiquitin-mediated proteasome system, all of which are major key nodes interlaced in the dual positive and negative feedback circuits, in order to maintain normal robust proteostasis.

## Discussion

After the NHB1 signal peptide of Nrf1 enables it to be anchored by a proper topology within the ER membranes, some portions of its other functional domains are translocated into the lumen, in which it is allowed for N-linked glycosylation by the luminal-resident active center of hetero-oligometic oligosaccharyltransferase (OST) (1,14). During retrotranslocation of Nrf1 glycoprotein into the cytoplasmic side of membranes, its subsequent deglycoprotein is further modified by ubiquitination and ensuing deubiquitination. All these modified proteins of Nrf1 are subjected to its selective topovectorial regulation in their juxtamembrane proteolytic processing to yield multiple distinct isoforms (15,32,33). Once deglycoprotein of Nrf1 by peptide:N-Glycanases (PNGase) is dislocated from the ER and transferred into the nucleus, it can function as a *bona fide* active form to regulate transcriptional expression of its cognate target genes (33,34). Similar results have also been obtained from the *Caenorhabditis elegans* homologue Skn-1 (35).

### Nrf1 is regulated by USP19 through its deubiquitination in close proximity to the ER

In close vicinity to the ER, Nrf1 is also subjected to ubiquitination by Hrd1 and other ubiquitin E3 ligases, before being targeted for the ubiquitin-mediated proteasomal degradation, particularly under normal conditions (12,14,18). However, it is worth noting that dislocation of Nrf1 from the ER membranes is largely unaffected by both its ubiquitination (15) and ensuing deubiquitination (in this study). Rather, it should also be noted that the selective proteolytic processing of Nrf1 appears to be regulated by its putative deubiquitination of Nrf1 by USP19, but not by USP14 or USP15, because our evidence has uncovered that this deubiquitinase possesses an ability to augment the stability of Nrf1 deglycoprotein and other processed proteins (Fig. 6D).

Herein, we found that two membrane-tethered proteins USP19 and Nrf1 are co-localized around and within the ER, such that this deubiquitinase can directly interact with the latter CNC-bZIP protein in close proximity to the ER membranes. The interaction of both proteins appears to depend on two regions of Nrf1 between amino acids 2-10 and/or 226-242. The former residues 2-10 of Nrf1 comprise the n-region of its ER-anchored NHB1 peptide, which resides in the cytoplasm, where it is allowed for potential interaction with the cytosolic ubiquitin-specific protease USP19. By contrast, the residues 226-242 of Nrf1 should be generally buried in the ER lumen and thus segregated from the cytosolically-resident USP19 (32,33). Thereby, it is inferred that only after this ETGE-containing region within Nrf1 will be dynamically moved out of membranes to enter the cytoplasmic side, it can enable the CNC-bZIP protein to interact with USP19. This notion is further corroborated by our findings that deletion of the n-region of ER-targeting signal in Nrf1^Δ2-10^ results in the resulting disappearance of deubiquitinated Nrf1 isoform, while Nrf1^Δ226-242^ gives rise to a constitutive emergence of such a similar deubiquitinated Nrf1 isoform.

Further evidence has also been presented that deubiquitination of the ER-resident Nrf1 by USP19 confers it to be emerging as a specific deubiquitinated isoform. The deubiquitinating effect of USP19 on Nrf1 (or its N298 region) is completely abolished by either USP19^C506S^ (a mutant of its enzymatic active center) or USP19^ΔTM^ (a deletion mutant of its C-terminal transmembrane domain anchored in the ER). These indicate that only the membrane-anchored USP19 gains accessibility to ER-associated Nrf1, before its deubiquitinating reaction with this protease. Moreover, another lines of supportive evidence have also been obtained from our experiments, unraveling that deubiquitination of Nrf1 by USP19 occurs only after p97-driven retrotranslocation of the luminal-resident CNC-bZIP glycoprotein into extra-ER compartments, in which it is further subjected to deglycosylation, deubiquitination and/or other proteolytic processing by distinct cytosolic enzymes. Such complex processing and modifications of Nrf1 allow it to express multiple protein isoforms, as determined by different electrophoresis systems (e.g., based on SDS-PAGE and LDS-NuPAGE gels) to show distinct lengths of Nrf1 protein isoforms [for its detailed descriptions, see the references (14,15)], one of which is its deubiquitinating isoform identified herein.

A similar USP19-deubiquitinating isoform of Nrf1 also seems to arise from expression of Nrf1^6×K/R^ alone, and this mutant deubiquitinating isoform is unaffected by this protease. This demonstrates that deubiquitination of Nrf1 by USP19 occurs at all six examined lysines with its NTD and AD1 regions (Fig. 4A). In-depth insights of distinct mutants and relevant recovery from Nrf1^6×K/R^ further revealed that the Lys^70^ is a key site for ubiquitination of Nrf1 and ensuing deubiquitination by USP19, whilst three other sites at Lys^169^, Lys^199^ and Lys^205^ within the N298 region of this CNC-bZIP protein are required for its ubiquitination and deubiquitination. However, such a couple of reversible modifications depends on the topological locations of all the four lysine sites (Lys^70^, Lys^169^, Lys^199^ and Lys^205^) within distinct contexts of Nrf1 or its mutants. This is due to the fact that Nrf1 ubiquitination and its deubiquitination by USP19 occur only after dynamic repositioning of these lysines from ER luminal side of membranes into the cytoplasmic side.

Notably, Nrf1^Δ2-10^ had been shown to give rise to a major strong glycoprotein, with little or none in its processed proteins (1,32). Similar results were also obtained from its N298^Δ2-10^ fusion protein (15), which is endowed as a major full-length protein of 80-kDa, but its proteolytic processing was substantially diminished by lack of the n-region of its NHB1 signal. As such, the remnant N-terminally-processed proteins of between 55-kDa and 12.5-kDa from N298^Δ2-10^ fusion mutant were also found to be almost unaffected by USP19. These observations demonstrate that the Lys^5/6^-containing n-region of NHB1 signal is required for a proper membrane-topology of this CNC-bZIP protein folding within and around the ER and its subsequent repositioning into the cytoplasmic side of membranes. Hence, it is inferable that putative Lys^5/6^ ubiquitination of Nrf1 by Hrd1 and its ensuing deubiquitination by USP19 are much likely to promote dynamic repartitioning of this CNC-bZIP protein into extra-ER subcellular compartments and its subsequent proteolytic processing by cytosolic proteasome, DDI1/2 and other proteases. This is also supported by our finding that loss of Lys^5/6^ in Nrf1^Δ2-10^ leads to constitutive disappearance of deubiquitinated Nrf1 isoform.

The effect of USP19 on Nrf1 to erase its conjugated ubiquitins renders this CNC-bZIP protein to circumvent the ubiquitin-led proteasomal degradation, such that abundances of its deglycoprotein and derivative proteins (arising from the selective proteolytic processing by DDI1/2 and/or other proteases) are enhanced by this deubiquitinatase. However, this manifests that the deubiquitinating effect of USP19 on Nrf1 processing is limited. As a matter of fact, the selective proteolytic processing of Nrf1 by cytosolic proteases (e.g., DDI1/2 (15,36,37)) is dictated by its intrinsic topologically-regulatory mechanism. This notion is established on the solid ground that distinct deletions of NHB1- and NHB2-adjoining peptides from Nrf1 or its N298 region result in complete abolishment or significant diminishment of its N-terminal DDI1/2-cleaved polypeptide with variations in an about 12.5-kDa molecular weight (Fig. S5). Such an objective fact demonstrates that no DDI-specific cleavage sites exist within Nrf1, and even if doing so, this is determined by topovectorial processes of this CNC-bZIP protein to be repositioned into the extra-ER side of membranes. This is further supported by another variation in the yield of other two N-terminally-cleaved polypeptides between 35-kDa and 55-kDa from the N298 fusion protein, in which their putative cleavage sites are deduced to be embodied within discrete peptide bonds of its AD1 domain. Within the N-terminal one-third of AD1, two DIDLID/DLG-adjoining elements (aa 125-170 & 171-186) are required for the stability of Nrf1 (and/or its N298) fusion proteins and its processing. The relevant mechanisms had been also elucidated in detail by (13,15,33).

Moreover, a not very satisfactory result obtained from *in vitro* enzymatic reactions (as shown in Fig. 1E) revealed the immunoprecipitates of Nrf1 could be only partially deubiquitinated by USP19 (implying that other deubiquitinating enzymes are involved in this biochemical process) and/or its proper enzymatic reaction may also be constrained only in the real place closer to the cytoplasmic side of the ER. This is inferable that the proper enzymatic reaction of USP19 to mediate deubiquitination of Nrf1 could require for certain unidentified coenzymes to work together. In fact, USP19 has many functions associated with the ER, including those involved in both ERAD (endoplasmic-reticulum-associated degradation) and MAPS (misfolding-associated protein secretion). This is fully consistent with the report by Hassink, *et al*, revealing that no cleavage activity of USP19 was determined upon deletion of its C-terminal TM domain (25). Thereby, the proper membrane-topology of USP19 attached to the ER plays an essential role in the functioning of this protease to mediate deubuquitination of Nrf1 in proper place. Such subcellular co-location of USP19 and Nrf1 around the ER compartments may also be beneficial to their functional combination with the other co-enzymatic factors.

### Nrf1 and Nrf2 have bidirectional inter-regulatory roles in co-targeting USP19 expression

Since USP19 enhances stability of deglycoprotein of Nrf1 and its processed proteins by escaping from ubiquintin proteasomal degradation, this is accompanied by an evident increase in the transcriptional activity of this CNC-bZIP factor to mediate expression of its target genes encoding proteasomal subunits. In turn, knockout of *USP19* leads to an obvious decrease in the proteasomal expression, so that the resulting ubiquitinated proteins are accumulated in *USP19*^−/−^ cells. As such, *USP19*^−/−^ cells still possess a modest resistance to cytotoxicity of the proteasomal inhibitor bortezomib, when compared to the equivalent of *Nrf1α*^−/−^ cells (Fig. S7C). This implicates only partial inactivation of Nrf1 to regulate the proteasomal gene expression in *USP19*^−/−^ cells, as evidenced by our further experiments. As a consequence, *USP19*^−/−^-derived tumor growth in xenograft model mice was modestly retarded, when compared with wild-type controls. This is attributable to the arrest of *USP19*^−/−^ cell cycle at its G0/G1 phase with a shortened S phase (in this study), whereas *Nrf1α*^−/−^ cell cycle is arrested at its G2/M phase (38,39). However, as *Nrf1α*^−/−^-derived tumor growth is significantly incremented, it is further deteriorated and metastasized to the liver and lung in xenograft mice (38-40). Such complete loss of Nrf1α-derived proteins results in an aberrant accumulation of Nrf2, as accompanied by inactivation of the tumor-repressor PTEN. By contrast, expression of Nrf2 at mRNA and protein levels appear to be unaffected by partial inactivation of Nrf1 in *USP19*^−/−^ cells. Altogether, these demonstrates that the remaining Nrf1’s function can still be exerted as a dominant tumor-repressor, particularly under Nrf2-unaffected conditions in *USP19*^−/−^ cells. Contrarily, hyper-active Nrf2 acts as a predominant functional rebel to become a cancer promoter in *Nrf1α*^−/−^ cells, but it is successfully confined by the remnant Nrf1 in *USP19*^−/−^ cells.

Within the regulatory feedback circuit, USP19 is positively regulated by Nrf2, but its transcriptional expression is only less or not promoted by accumulated Nrf2 in *Nrf1α*^−/−^ cells. This implies that positive regulation of USP19 by Nrf2 appears to be suppressed by a not-yet-identified mechanism in *Nrf1α*^−/−^ cells. In other words, Nrf1α-derived factors also indirectly contribute to positive regulation of USP19, albeit this protease is also subjected to a negative feedback loop governed by Nrf1-target proteasomal genes (Fig. 6D). In fact, ectopic over-expression of Nrf1, like Nrf2, leads to an increase in transcription activity of *USP19*-driven reporter by non-ARE consensus sites. Such striking disparity in between effects of Nrf1 and Nrf2 on the endogenous *USP19* and its promoter-driven reporter (i.e., *USP19-Luc*) genes indicates that differential transcriptional regulation of their expression could also depend on their genomic contexts of distinct topological settings.

In summary, this work provides a holistic approach together with the reductionist’s insights into those key nodes (as illustrated in Fig. 6D), which are topologically organized together within a complex hierarchical regulatory network, in order to maintain and perpetuate the steady state of distinct cellular homeostasis and subcellular organelle integrity. The robustness and plasticity of the homeostasis (including proteostasis) are also dependent on its negative feedback loop, in terms of system dynamics to persist the functional stability of this complex regulatory network. For this reason, to gain a holistic view of Nrf1 along with Nrf2 and USP19 in their inter-regulatory network, we have comprehensively investigated distinct aspects of them from several distinct angles (at once when we can do so as possible) by different experimental settings of this study done in different conditions, but not only by favoring a piecemeal approach usually focused on Nrf1 regulation by USP19. Consequently, our evidence has been presented in this study, demonstrating that USP19 serves as a novel mechanistic modulator of Nrf1, but not Nrf2, allowing the former CNC-bZIP protein to be rescued from the ubiquitin-directed ER-associated degradation. Conversely, differential transcriptional expression of this protease USP19 is also regulated by Nrf2, as well by Nrf1, at distinct layers within bidirectional regulatory feedback circuits. Besides USP19, most of distinct deubiquitinase family members are also, to greater or less degrees, regulated by Nrf1 or Nrf2 alone or both factors.

## Materials and Methods

### Cell lines, Culture and Transfection

Knockout cell lines of *USP19*^−/−^ were here created by CRISPR/Cas9-mediated gene manipulation on the basis of HepG2. Briefly, single guide RNAs targeting exons 3 of the *USP19* gene (#1, 5′-AGGAAGCCCGAACCAGAAGCGG-3′; #2, 5′-ATGCATCAAACCGTGAGCAGCGG-3′) were cloned into a pCAG-T7-cas9+gRNA-pgk-Puro-T2A-GFP vector (Viewsolid Biotech, Beijing, China). The plasmid was then transfected into HepG2 cells with Lipofectamine 3000 (Invitrogen, Carlsbad, CA), followed by selecting positive cells with puromycin (2.5 µg/ml). The anti-drug monoclonal cells (all of which had been originated from a single cell) were identified by the genomic DNA locus-specific sequencing. The *Nrf1*^−/−^ cells were established by TALANs-led genome editing (38). The *Nrf2*^−/−^ cells were constructed by CRISPR/Cas9-editing system (39). These cell lines were maintained in DMEM supplemented with 5 mM glutamine, 10% (v/v) foetal bovine serum (FBS), 100 units/mL of either of penicillin and streptomycin, in the 37°C incubator with 5% CO_2_. In addition, some of cell lines were transfected for 8 h with the indicated constructs mixed with the Lipofectamine^®^3000 agent in the Opti-MEM (gibca, Waltham, MA, USA). The cells were then allowed for recovery from transfection in a fresh complete medium for 24 h, before the other experiments were conducted.

### Expression constructs

Five expression constructs for human USP14, USP15, USP19, USP30 and USP48 were here created by inserting their full-length cDNA sequences into the p3xFlag-CMV-14 vector. Additional two mutants USP19^ΔTM^ and USP19^C506S^ were also made. The former USP19^ΔTM^ was constructed by deleting a cDNA fragment encoding the transmembrane-relevant residues 1393-1413 of USP19, while the latter USP19^C506S^ mutant was yielded by replacing its original cysteine at position 506 with a serine residue. Notably, 4202-bp of the *USP19* promoter and its ARE-indicated mutants were cloned into the KpnI/HindIII site of the PGL3-Basic vector to yield wild-type *USP19-Luc* and its corresponding mutant reporters. In addition, two expression constructs for human Nrf1 and Nrf2 were made by inserting the full-length cDNA sequences into the KpnI/XbaI site of pcDNA3.1/V5His B, as described previously by (1,2). A set of mutants of those putative ubiquitinated lysine sites and relevant recovery plasmids from its mutation was created on the base of a Nrf1 expression construct. Another set of N298 fusion protein expression construct and its deletion mutants was also constructed by inserting the N-terminal first 298-aa of Nrf1 into peGFP-N2 vector. Of note, the N-terminus of N298 was tagged by attaching the V5 epitope, as described elsewhere (15).

### Real-time qPCR analysis of mRNA expression

Approximately 2.5 µg of total RNAs (obtained by using an RNeasy mini-RNAsimple Kit, Tiangen Biotech, Beijing, China) were added in a reverse-transcriptase reaction to generate the first strand of cDNA (by using the Revert Aid First Strand Synthesis Kit, Thermo, Waltham, MA, USA). The synthesized cDNA was served as the template for qPCR, in the GoTaq^®^qPCR Master Mix (Promega, Madison, WI, USA), before being deactivated at 95°C for 10 min, and then amplified by 40 reaction cycles of being annealed at 95°C for 15 s and then extended at 60°C for 30 s. The final melting curve was validated to examine the amplification quality. Of note, the mRNA expression level of β-actin served as an optimal internal standard control, relative to other mRNA expression levels presented as fold-changes. All the forward and reverse primers of those indicated genes were shown in Table S1.

### Western blotting and pulse-chase experiments

Experimental cells were harvested in a denatured lysis buffer (0.5% SDS, 0.04 mol/L DTT, pH 7.5, containing 1 tablet of cOmplete protease inhibitor EASYpacks in 10 ml of this buffer). The total lysates were further denatured by boiling at 100°C for 10 min, sonicated sufficiently, and diluted with 3× loading buffer (187.5 mmol/L Tris-HCl, pH 6.8, 6% SDS, 30% Glycerol, 150 mmol/L DTT, 0.3% Bromphenol Blue), before being re-boiled at 100°C for 5 min. Thereafter, equal amounts of protein extracts were subjected to separation by either SDS-PAGE containing 8% or 10% polyacrylamide or LDS-NuPAGE containing 4-12% polyacrylamide, in two distinct pH running buffers, before being transferred to polyvinylidene fluoride (PVDF) membranes (Millipore, Billerica, MA, USA). The protein-transferred membranes were blocked for 1 h by 5% fat-free milk (w/v) resolved in PBS containing 0.1% Tween 20 (v/v) and then incubated with one of indicated primary antibodies for 2 h at room temperature or overnight at 4°C, followed by washing steps. The antibody-incubated membranes were visualized by 2 h of re-incubation with an appropriate horseradish peroxidase-conjugated secondary antibodies, followed by enhanced chemiluminescence. To estimate the protein stability, its half-life was calculated by cycloheximide (CHX)-pulse chase experiments of distinct cell lines that had been transfected or not transfected with the indicated expression constructs and treated with 50 µg/ml CHX for an indicated time, followed by immunoblotting of these cell lysates.

### *In vitro* deglycosylation reactions

Experimental cells were harvested in a denatured lysis buffer and denatured by boiling at 100°C for 10 min as above. The heat-denatured samples were incubated at 37 °C for 1 h with 500 units of Endo H (endoglycosidase H; New England Biolabs, Ipswich, MA) in a final volume of 20 µl containing 1 × GlycoBuffer. The reaction products were subjected to protein separation by SDS-PAGE or LDS-NuPAGE and visualization by Western blotting.

### Luciferase Reporter Assay

Equal numbers (1.0 × 10^5^) of COS1 cells were allowed for growth in each well of 12-well plates. After reaching 80% confluence, the cells were co-transfected for 8 h with an indicated luciferase plasmid alone or together with one of expression constructs in the Lipofectamine 3000 mixture, in which the Renilla-expressing pRL-TK plasmid served as an internal control for transfection efficiency. The cells were allowed for a recovery from transfection to be cultured in a fresh complete medium for 24 h, before the luciferase activity was measured by the dual-reporter assay (Promega, Madison, WI, USA).

### Subcellular fractionation with the ER isolation

Equal numbers (1 × 10^6^) of distinct cell lines were allowed for growth in each of 6-cm dishes for 24 h before being harvested by incubation with 1 mL of an ice-cold nuclei EZ lysis buffer (sigma, San Francisco, CA, USA). The total lysates were subjected to subcellular fractionation by centrifuging at 500×*g* for 5 min at 4°C. The supernatants were collected as a non-nuclear cytoplasmic fraction, whilst the sediment fraction were further washed with the above lysis buffer for two times, each of which 0.5 mL of the nuclei EZ lysis buffer was added into the sediment fraction. Lastly, the nuclear pellets were obtained by centrifuging at 500×*g* for 5 min at 4°C, whereas the ER fractions were further obtained by centrifuging the extra-nuclear supernatant at 10000×*g* for 5 min at 4°C. Subsequently, these fractions were further evaluated by Western blotting.

### Co-immunoprecipitation with ubiquitination and deubiquitination analysis

Experimental cells were lysed in 500 µL of the NP-40 lysis buffer (20 mM Tris-HCl, pH 7.4∼7.5, 150 mM NaCl, 1 mM EDTA, 1% NP-40, and 1 tablet of cOmplete protease inhibitor EASYpacks resolved in 10 ml of this buffer). The lysates were centrifuged at 12,000 rpm (=12396×*g*) for 10 min at 4°C to obtain the clear supernatants. Subsequently, the supernatants was subject to immunoprecipitation by being incubated with 1.2 μg of specific antibody overnight, and re-incubated with 20 μL of Protein A/G PLUS-Agarose (santa cruz, Santa Cruz, CA, USA) at 4°C for 2 h. After the beads were washed for three times, each with 1 ml of the above lysis buffer containing a 500 mM concentration of NaCl, the resulting immunoprecipitates were resolved by SDS-PAGE gels and visualized by immunoblotting with indicated antibodies. For analysis of ubiquitination and deubiquitination status, the ubiquitin-expressing cell lysates were analyzed by Western blotting with its specific antibody or by immunoprecipitation with its HA-tag antibody. For analysis of deubiquitination of Nrf1 by USP19, Nrf1-expressing lysates were allowed for *in vitro* reactions with USP19 immunoprecipitates at 37°C for different times. Then, the electrophoretic mobility of Nrf1 was then determined by Western blotting with V5 antibody.

### Immunocytochemistry with confocal microscopy

Experimental cells (3 × 10^5^), which had been allowed for overnight growth on a cover glass being placed in each well of 6-well plates, were co-transfected for 8 h with two distinct expression constructs for USP19 plus Nrf1 or its mutant and then allowed for 24 h recovery from transfection in the fresh complete medium. Thereafter, these cells were fixed for 15 min with 4% paraformaldehyde in PBS buffer and then permeabilized for additional 10 min with 0.1% Triton X-100 in PBS. Subsequently, the fixed cells were subjected to immunocytochemistry by incubation with the primary antibodies against the V5-epitoped tag or USP19 (each with a dilution of 1:200) for 2 h and additional incubation with either Alexa Fluor 488-conjugeted Goat Anti-Mouse IgG (H+L) or TRITC-conjugeted Goat anti-Rabbit IgG (H+L) (both obtained from ZSGB-BIO, Beijing, China) for 2 h at room temperature in the dark, followed by an additional 5-min staining of DNA with DAPI (Beyotime, Beijing, China). The results of distinct fluorescence images were achieved by confocal microscopy.

### Subcutaneous tumor xenografts in nude mice with pathohistological analysis

Mouse xenograft models were made by subcutaneous heterotransplantation of wild-type human hepatoma cells or its derived *USP19*^−/−^ cells in nude mice. Briefly, equal amounts of experimental cells (1 × 10^7^) growing in the exponential phase were suspended in 0.1 ml of serum-free DMEM and then inoculated subcutaneously at a single site in the right upper back region of male nude mice (BALB/C nu/nu, 4–6 weeks, 18 g). The procedure of injection into all the mice was completed within 30 min. Subsequently, the formation of these murine subcutaneous tumor xenografts was observed successively until all these mice were sacrificed. The transplanted tumors were excised immediately after they were executed. The tumor sizes were also calculated by a standard formulate (i.e. V = ab^2^/2), as shown graphically (n= 4 per group). Furthermore, the tumor tissues were immersed in 4% paraformaldehyde overnight and then transferred to 70% ethanol. The fixed tumor tissues were dehydrated by a serial gradient of, alcohol, embedded in paraffin wax blocks, and sectioned into a series of 5-μm-thick slides. Before pathohistological staining, the sections were de-waxed in xylene and rehydrated by progressively-decreased concentrations of ethanol. Subsequently, these tissue sections were stained routinely by hematoxylin and eosin (H&E) and then visualized by microscopy. In addition, it should also be noted that these nude mice were purchased from the Laboratory Animal Center of Chongqing Medical University (with a certificate SCXK (YU) 2007-0001). They were maintained under the optimal conditions for hygiene, temperature and photoperiods (12L:12D), and allowed *ad libitum* to food and water, according to the institutional guidelines for the care and use of laboratory animals. All the experimental procedures were approved by the Ethics Committee of Chongqing medical University.

### Cell viability analysis

All three cell lines of wild-type HepG2, *Nrf1α*^−/−^ and *USP19*^−/−^ were cultured for 24 h in 96-well plates. After reaching 70% of their confluence, they were allowed for growth in fresh media containing different concentrations of *bortezomib* (BTZ) (at 0, 0.01, 0.1, 1, 10 or 100 µmol/L) for 24h, which was dissolved in DMSO (dimethyl sulfoxide; 0.1% of this solvent was herein used as a vehicle control). The cell viability was then evaluated by using an MTT-based cell proliferation and cytotoxicity assay kit (Beyotime, Shanghai, China).

### Flow cytometry analysis of cell cycle and apoptosis

Experimental cells (6 × 10^5^) were allowed for growth in 60-mm cell culture plate for 48 h and synchronization by 12-h starvation in a serum-free medium, before being treated with 10 μmol/L BrdU for 12 h. The cells were fixed for 15 min with 100 μL of BD Cytofix/Cytoperm buffer (with a mixture of both the fixative paraformaldehyde and the detergent saponin) at room temperature and then permeabilized for 10 min in 100 μL BD Cytoperm permeabilization buffer plus (with fetal bovine serum served as a staining enhancer) on ice. Thereafter, the cells were treated for 1 h at 37 °C with 100 μL of DNase (at a concentration of 300 μg/mL in PBS) to allow for exposure to BrdU incorporation, before being stained with fluorescein isothiocyanate conjugated anti-BrdU antibody for 60 min at room temperature. Subsequently, the cells were suspended in 20 μL of a 7-amino-actinomycin D solution for 20-min DNA staining and re-suspended in 0.5 mL of another staining buffer (1 × DPBS containing 0.09% sodium azide and 3% heat-inactivated FBS), prior to flow cytometry analysis of cell cycles. Furthermore, additional fractions of cells were allowed for 48-h growth in 60-mm cell culture plate and subjected to apoptosis analysis by flow cytometry. The cells were pelleted by centrifuging at 1000×*g* for 5 min and washed by PBS for three times, before being incubated for 15 min with 5 μL of Annexin V-FITC and 10 μL of propidium iodide (PI) in 195 μL of the binding buffer. The results were analyzed by the FlowJo 7.6.1 software (FlowJo, Ashland, OR, USA) before being presented.

### Statistical analysis

Statistical significance of changes in reporter activity and other gene expression was determined using either the Student’s t-test or Multiple Analysis of Variations (MANOVA). The resulting data are shown as a fold change (mean ± S.D), each of which represents at least 3 independent experiments that were each performed triplicate.

## Author contributions

S.H. performed most experiments with help of L.Q. and M.W., and collected all the relevant data, except that Y.X. did a few of experiments as shown in Figure 1. S.H. also made draft of this manuscript with most figures and supplemental information. Y.Z. designed and supervised this study, analyzed all the data, helped to prepare all figures with cartoons, wrote and revised the paper.

## Acknowledgments

We are greatly thankful to Dr. Yonggang Ren (North Sichuan Medical College, Sichuan, China) for his involvement in establishing the indicated cell lines used in this study. We also thank to Mr. Ze Zheng and other members of Prof. Zhang’s laboratory (at Chongqing University, China) for giving their invaluable help with this work. Notably, this study was funded by the National Natural Science Foundation of China (NSFC, with a key program 91429305 and additional two projects 81872336 and 82073079) awarded to Prof. Yiguo Zhang.

## References

1. Zhang, Y., Lucocq, J. M., Yamamoto, M., and Hayes, J. D. (2007) The NHB1 (N-terminal homology box 1) sequence in transcription factor Nrf1 is required to anchor it to the endoplasmic reticulum and also to enable its asparagine-glycosylation. The Biochemical journal 408, 161–172

2. Zhang, Y., Crouch, D. H., Yamamoto, M., and Hayes, J. D. (2006) Negative regulation of the Nrf1 transcription factor by its N-terminal domain is independent of Keap1: Nrf1, but not Nrf2, is targeted to the endoplasmic reticulum. The Biochemical journal 399, 373–385

3. Sykiotis, G. P., and Bohmann, D. (2010) Stress-activated cap’n’collar transcription factors in aging and human disease. Science signaling 3, re3

4. Zhang, Y., and Xiang, Y. (2016) Molecular and cellular basis for the unique functioning of Nrf1, an indispensable transcription factor for maintaining cell homoeostasis and organ integrity. The Biochemical journal 473, 961–1000

5. Zhu, Y. P., Wang, M., Xiang, Y., Qiu, L., Hu, S., Zhang, Z., Mattjus, P., Zhu, X., and Zhang, Y. (2018) Nach Is a Novel Subgroup at an Early Evolutionary Stage of the CNC-bZIP Subfamily Transcription Factors from the Marine Bacteria to Humans. International journal of molecular sciences 19, 1–26

6. Lynn, D. A., and Curran, S. P. (2015) The SKN-1 hunger games: May the odds be ever in your favor. Worm 4, e1078959

7. Yamamoto, M., Kensler, T. W., and Motohashi, H. (2018) The KEAP1-NRF2 System: a Thiol-Based Sensor-Effector Apparatus for Maintaining Redox Homeostasis. Physiological reviews 98, 1169–1203

8. Gegotek, A., and Skrzydlewska, E. (2015) CNC proteins in physiology and pathology. Postepy Hig Med Dosw (Online) 69, 729–743

9. Bugno, M., Daniel, M., Chepelev, N. L., and Willmore, W. G. (2015) Changing gears in Nrf1 research, from mechanisms of regulation to its role in disease and prevention. Biochim Biophys Acta 1849, 1260–1276

10. Hayes, J. D., Dinkova-Kostova, A. T., and Tew, K. D. (2020) Oxidative Stress in Cancer. Cancer cell 38, 167–197

11. Zhu, Y. P., Zheng, Z., Hu, S., Ru, X., Fan, Z., Qiu, L., and Zhang, Y. (2019) Unification of Opposites between Two Antioxidant Transcription Factors Nrf1 and Nrf2 in Mediating Distinct Cellular Responses to the Endoplasmic Reticulum Stressor Tunicamycin. Antioxidants 9, 4

12. Steffen, J., Seeger, M., Koch, A., and Kruger, E. (2010) Proteasomal degradation is transcriptionally controlled by TCF11 via an ERAD-dependent feedback loop. Molecular cell 40, 147–158

13. Widenmaier, S. B., Snyder, N. A., Nguyen, T. B., Arduini, A., Lee, G. Y., Arruda, A. P., Saksi, J., Bartelt, A., and Hotamisligil, G. S. (2017) NRF1 Is an ER Membrane Sensor that Is Central to Cholesterol Homeostasis. Cell 171, 1094–1109

14. Xiang, Y., Wang, M., Hu, S., Qiu, L., Yang, F., Zhang, Z., Yu, S., Pi, J., and Zhang, Y. (2018) Mechanisms controlling the multistage post-translational processing of endogenous Nrf1alpha/TCF11 proteins to yield distinct isoforms within the coupled positive and negative feedback circuits. Toxicology and applied pharmacology 360, 212–235

15. Xiang, Y., Halin, J., Fan, Z., Hu, S., Wang, M., Qiu, L., Zhang, Z., Mattjus, P., and Zhang, Y. (2018) Topovectorial mechanisms control the juxtamembrane proteolytic processing of Nrf1 to remove its N-terminal polypeptides during maturation of the CNC-bZIP factor. Toxicology and applied pharmacology 360, 160–184

16. Sha, Z., and Goldberg, A. L. (2014) Proteasome-mediated processing of Nrf1 is essential for coordinate induction of all proteasome subunits and p97. Current biology: CB 24, 1573–1583

17. Hamazaki, J., and Murata, S. (2020) ER-Resident Transcription Factor Nrf1 Regulates Proteasome Expression and Beyond. International journal of molecular sciences 21, 3683

18. Tsuchiya, Y., Morita, T., Kim, M., Iemura, S., Natsume, T., Yamamoto, M., and Kobayashi, A. (2011) Dual regulation of the transcriptional activity of Nrf1 by beta-TrCP-and Hrd1-dependent degradation mechanisms. Molecular and cellular biology 31, 4500–4512

19. Baldridge, R. D., and Rapoport, T. A. (2016) Autoubiquitination of the Hrd1 Ligase Triggers Protein Retrotranslocation in ERAD. Cell 166, 394–407

20. Hwang, J., Walczak, C. P., Shaler, T. A., Olzmann, J. A., Zhang, L., Elias, J. E., and Kopito, R. R. (2017) Characterization of protein complexes of the endoplasmic reticulum-associated degradation E3 ubiquitin ligase Hrd1. The Journal of biological chemistry 292, 9104–9116

21. Ventii, K. H., and Wilkinson, K. D. (2008) Protein partners of deubiquitinating enzymes. The Biochemical journal 414, 161–175

22. Villeneuve, N. F., Tian, W., Wu, T., Sun, Z., Lau, A., Chapman, E., Fang, D., and Zhang, D. D. (2013) USP15 negatively regulates Nrf2 through deubiquitination of Keap1. Molecular cell 51, 68–79

23. Fukagai, K., Waku, T., Chowdhury, A. M., Kubo, K., Matsumoto, M., Kato, H., Natsume, T., Tsuruta, F., Chiba, T., Taniguchi, H., and Kobayashi, A. (2016) USP15 stabilizes the transcription factor Nrf1 in the nucleus, promoting the proteasome gene expression. Biochemical and biophysical research communications 478, 363–370

24. Taniguchi, H., Okamuro, S., Koji, M., Waku, T., Kubo, K., Hatanaka, A., Sun, Y., Chowdhury, A. M., Fukamizu, A., and Kobayashi, A. (2017) Possible roles of the transcription factor Nrf1 (NFE2L1) in neural homeostasis by regulating the gene expression of deubiquitinating enzymes. Biochemical and biophysical research communications 484, 176–183

25. Hassink, G. C., Zhao, B., Sompallae, R., Altun, M., Gastaldello, S., Zinin, N. V., Masucci, M. G., and Lindsten, K. (2009) The ER-resident ubiquitin-specific protease 19 participates in the UPR and rescues ERAD substrates. EMBO reports 10, 755–761

26. Xu, Y., Cui, L., Dibello, A., Wang, L., Lee, J., Saidi, L., Lee, J. G., and Ye, Y. (2018) DNAJC5 facilitates USP19-dependent unconventional secretion of misfolded cytosolic proteins. Cell discovery 4, 11

27. Lei, C. Q., Wu, X., Zhong, X., Jiang, L., Zhong, B., and Shu, H. B. (2019) USP19 Inhibits TNF-alpha-and IL-1beta-Triggered NF-kappaB Activation by Deubiquitinating TAK1. Journal of immunology 203, 259–268

28. Miao, R., Lu, Y., He, X., Liu, X., Chen, Z., and Wang, J. (2020) Ubiquitin-specific protease 19 blunts pathological cardiac hypertrophy via inhibition of the TAK1-dependent pathway. Journal of cellular and molecular medicine 00, 1–12

29. Eletr, Z. M., and Wilkinson, K. D. (2014) Regulation of proteolysis by human deubiquitinating enzymes. Biochim Biophys Acta 1843, 114–128

30. Coyne, E. S., and Wing, S. S. (2016) The business of deubiquitination - location, location, location. F1000Research 5, 163

31. Clague, M. J., Urbe, S., and Komander, D. (2019) Breaking the chains: deubiquitylating enzyme specificity begets function. Nature reviews. Molecular cell biology 20, 338–352

32. Zhang, Y., Li, S., Xiang, Y., Qiu, L., Zhao, H., and Hayes, J. D. (2015) The selective post-translational processing of transcription factor Nrf1 yields distinct isoforms that dictate its ability to differentially regulate gene expression. Scientific reports 5, 12983

33. Zhang, Y., Ren, Y., Li, S., and Hayes, J. D. (2014) Transcription factor Nrf1 is topologically repartitioned across membranes to enable target gene transactivation through its acidic glucose-responsive domains. PloS one 9, e93458

34. Zhang, Y., Lucocq, J. M., and Hayes, J. D. (2009) The Nrf1 CNC/bZIP protein is a nuclear envelope-bound transcription factor that is activated by t-butyl hydroquinone but not by endoplasmic reticulum stressors. The Biochemical journal 418, 293–310

35. Lehrbach, N. J., Breen, P. C., and Ruvkun, G. (2019) Protein Sequence Editing of SKN-1A/Nrf1 by Peptide:N-Glycanase Controls Proteasome Gene Expression. Cell 177, 737–750

36. Lehrbach, N. J., and Ruvkun, G. (2016) Proteasome dysfunction triggers activation of SKN-1A/Nrf1 by the aspartic protease DDI-1. eLife 5, e17721

37. Koizumi, S., Irie, T., Hirayama, S., Sakurai, Y., Yashiroda, H., Naguro, I., Ichijo, H., Hamazaki, J., and Murata, S. (2016) The aspartyl protease DDI2 activates Nrf1 to compensate for proteasome dysfunction. eLife 5, e18357

38. Ren, Y., Qiu, L., Lü, F., Ru, X., Li, S., Xiang, Y., Yu, S., and Zhang, Y. (2016) TALENs-directed knockout of the full-length transcription factor Nrf1a that represses malignant behaviour of human hepatocellular carcinoma (HepG2) cells Scientific reports 6, 23775

39. Qiu, L., Wang, M., Hu, S., Ru, X., Ren, Y., Zhang, Z., Yu, S., and Zhang, Y. (2018) Oncogenic Activation of Nrf2, Though as a Master Antioxidant Transcription Factor, Liberated by Specific Knockout of the Full-Length Nrf1alpha that Acts as a Dominant Tumor Repressor. Cancers 10, 520

40. Chen, J., Wang, M., Xiang, Y., Ru, X., Ren, Y., Liu, X., Qiu, L., and Zhang, Y. (2020) Nrf1 Is Endowed with a Dominant Tumor-Repressing Effect onto the Wnt/beta-Catenin-Dependent and Wnt/beta-Catenin-Independent Signaling Networks in the Human Liver Cancer. Oxidative medicine and cellular longevity 2020, 5138539

