## Supplemental Figures for "Activation of the membrane-bound Nrf1 transcription factor by USP19, a ubiquitin-specific protease C-terminally tail-anchored in the endoplasmic reticulum"

**Supplementary Materials** included the relevant experimental data as shown in eight figures below, in addition to Table1, showing a list of all key reagents and resources used in this work.

#### Supplementary Figure S1

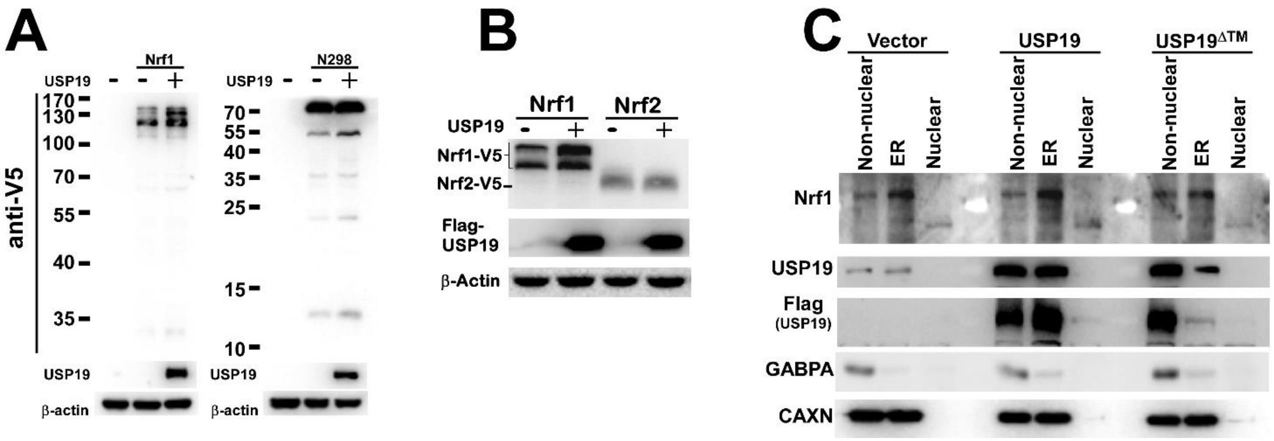


**Figure S1. The membrane-tethered USP19 exerts its enzymatic activity to mediate deubiquitination of Nrf1, but not Nrf2.**

(A) COS-1 cells had been co-transfected with distinct combinations of expression constructs for Nrf1-V5 (+), V5-N298-eGFP (+) or an empty vector (–), together with USP19-Flag (+) or the empty vector without this protease (–), and allowed for 24-h recovery from transfection, before the cells were harvested in denatured lysis buffer, then visualized by Western blotting with antibodies against V5 or USP19 epitopes.

(B) COS-1 cells co-expressing of either Nrf1-V5 or Nrf2-V5, together with USP19 (+), or not with this protease (–), were evaluated by Western blotting with V5 or Flag antibodies.

(C) HepG2 cells had been transfected with expression constructs for USP19, mutants USP19$^{\Delta TM}$ or an empty vector without this protease and allowed for 24-h recovery from transfection, before the cells were harvested. Thereafter, subcellular fractionation was conducted to isolate distinct fraction of the nucleus, and the endoplasmic reticulum, and extra-ER non-nuclear supernatants (i.e. cytoplasmic fractions), respectively, and visualized by immunoblotting with antibodies against Nrf1, USP19 or Flag epitopes.

### Supplementary Figure S2

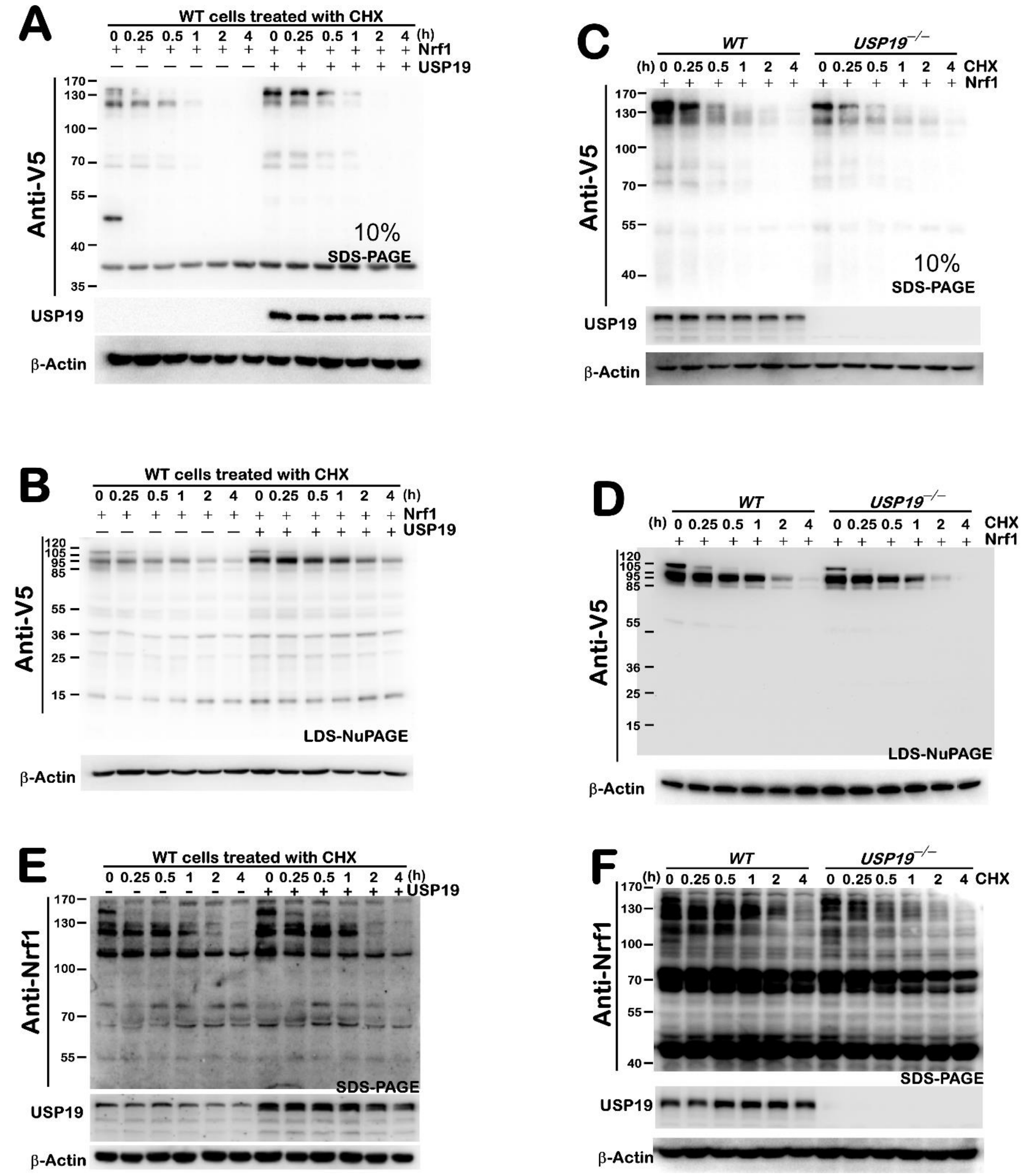


**Figure S2. USP19 has a marked effect on the stability of Nrf1 and its half-life.**

(A, B) COS-1 cells had been transfected with an expression construct for Nrf1 alone or plus USP19-Flag and allowed for 24-h recovery from transfection, before the cells were treated with CHX for indicated lengths of time and then harvested in denatured lysis buffer. Subsequently, the cell lyastes were separated by either SDS-PAGE gels containing 10% polyacrylamide (A) or LDS-NuPAGE gels containing 4-12% polyacrylamide (B), and visualized by immunoblotting with antibodies against V5 or Flag epitopes. The images cropped from the same gels are shown in the main Figure 2A.

(C,D) Total lysates of *WT* (HepG2) and *USP19*$^{-/-}$ cells, that had been allowed for ectopic over-expression of Nrf1-V5 and treated with CHX for indicated times, were resolved by 8% SDS-PAGE gels (C) or 4-12% LDS-NuPAGE gels (D), and visualized by Western blotting as described above. The images cropped from the same gels are shown in Figure 2C.

(E,F) CHX-pulse chase experiments of *WT* cells expressing USP19 or not (E), plus *USP19*$^{-/-}$ cells (F) were carried out as described above. The relevant images cropped from the same gels are shown in the main Figure 2E & 2G, respectively.

#### Supplementary Figure S3

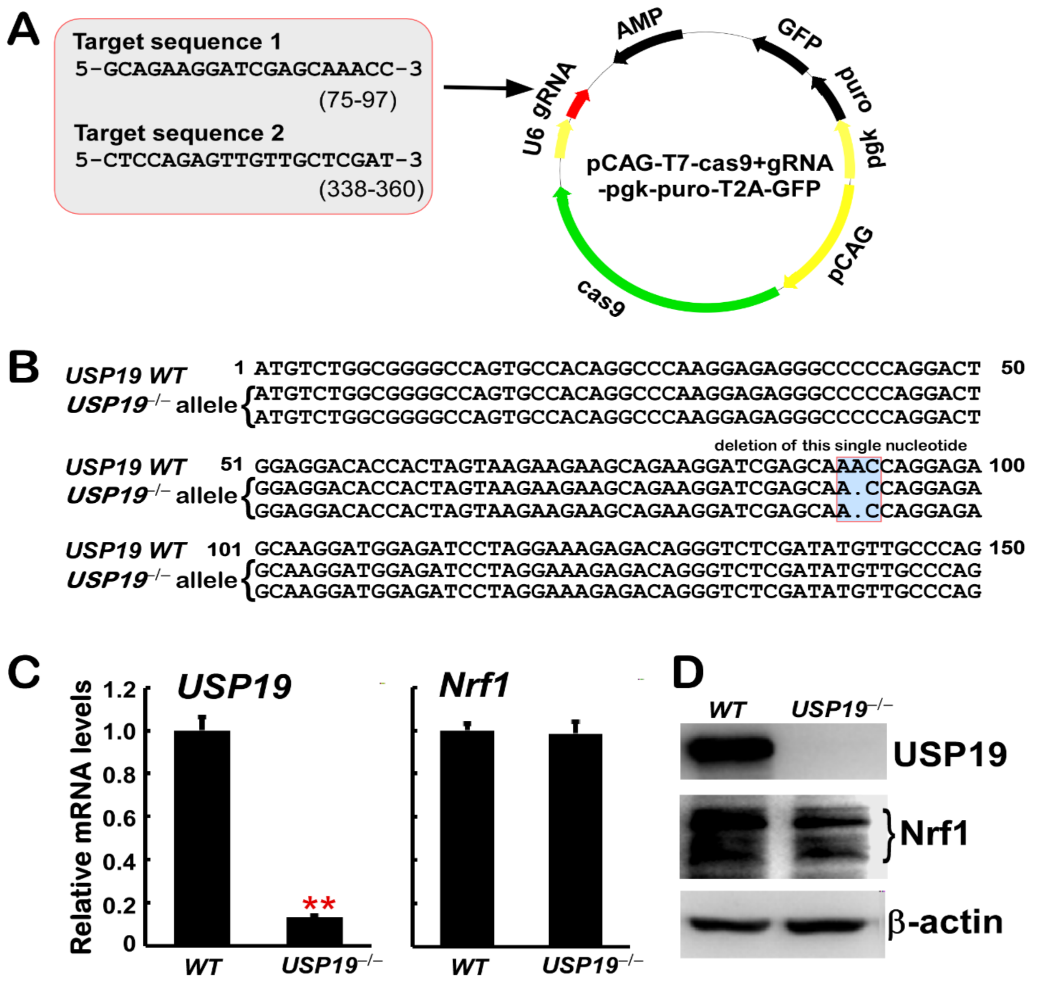


**Figure S3. Identification of *USP19*$^{-/-}$ cells by genomic locus sequencing, real-time qPCR and Western blotting.**

(A) A profile for CRISPR/CAS9-directed genome editing through two site-specific guide sequences targeting *USP19*.

(B) An alignment of site-specific genomic DNA sequences from *WT* and *USP19*$^{-/-}$ mutant alleles. The latter mutants contain a single nucleotide deletion within the 31$^{th}$ residue codon.

(C) Relative mRNA expression levels of *USP19* and *Nrf1* in *WT* and *USP19*$^{-/-}$ cells were determined by real-time qPCR. The resulting data are shown as fold changes (mean ± SEM, n= 3 × 3) with significant decreases (** $p$ < 0.001) as compared with the relevant controls.

(D) The above cell lysates were also subjected to Western blotting analysis of both protein levels of USP19 and Nrf1 in *WT* and *USP19*$^{-/-}$ cells, as described for the legend of Fig. S1.

### Supplementary Figure S4

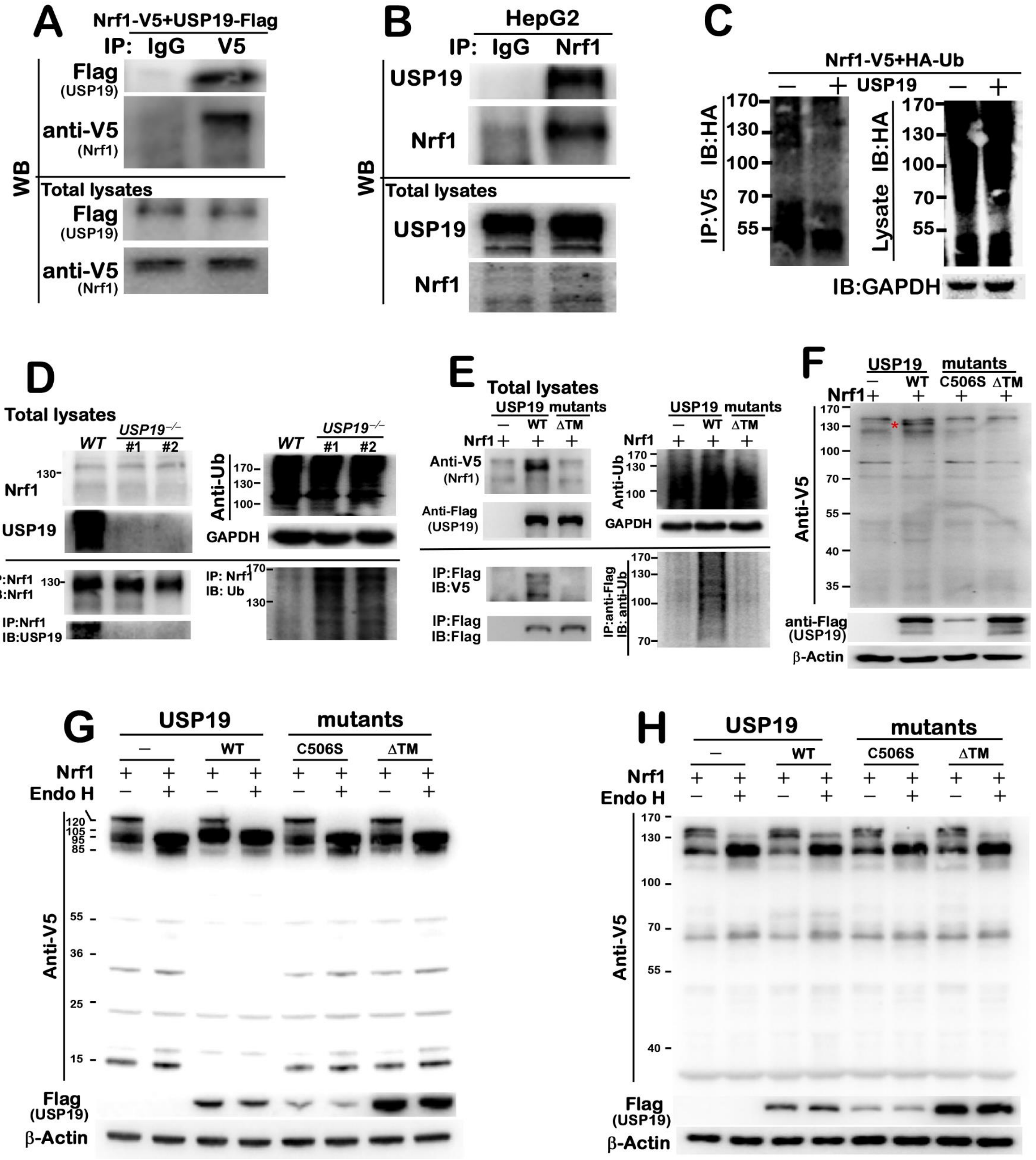


**Figure S4. Membrane-tethered USP19 exerts its enzymatic activity to mediate deubiquitination of Nrf1**

(A) Immunoprecipitates (IP) of COS-1 cells co-expressing USP19-flag and Nrf1-V5 recognized by two antibodies against IgG or V5 tag were determined by Western blotting (WB) with anti-Flag or -V5 antibodies, respectively (*upper two panels*). Besides, the inputs of whole-cell lysates were examined in parallel experiments (*lower two panels*).

(B) Immunoprecipitates (IP) containing endogenous proteins of HepG2 cells by antibodies against IgG or Nrf1 were analyzed by Western blotting (WB) with anti-USP19 or -Nrf1 antibodies, respectively (*upper two panels*). The inputs of whole-cell lysates were also subjected to Western blotting analysis with antibodies against USP19 or Nrf1 (*lower two panels*).

(C) Another anti-V5 immunoprecipitates (IP) of COS-1 cells co-expressing of Nrf1-V5 and HA-Ub, plus USP19 (+) or an empty vector without this protease (–), were visualized by anti-HA immunoblotting (IB) (*left panel*). The whole-cell lysates were also examined in the parallel experiments (*right panels*).

(D) Anti-Nrf1 immunoprecipitates of wild-type (*WT*) HepG2 and *USP19*$^{-/-}$ cell lines were examined by immunoblotting with antibodies against USP19, Nrf1 or Ub epitopes (*lower panels underlined*). The inputs of whole-cell lysates were also examined in the meantime (*upper panels*).

(E) Anti-Flag immunoprecipitates (IP) of COS-1 cells co-expressing of Nrf1-V5 and HA-Ub, together with USP19, its mutant USP19$^{\Delta TM}$ or an empty vector (as a negative control), were visualized by immunoblotting (IB) with antibodies against Ub, V5 or Flag (*lower panels underlined*)). The whole-cell lysates were examined in the parallel experiments (*upper panels*).

(F) Total lysates of COS-1 cells co-transfected with an expression construct for Nrf1 alone or plus USP19 (WT), its mutants USP19$^{C506S}$ or USP19$^{\Delta TM}$ were determined by Western blotting with antibodies against Flag or V5 tag, respectively.

(G, H) Total lysates of COS-1 cells was carried out as described above, and then allowed for *in vitro* deglycosylation reactions with Endoglycosidase H (Endo H) (+) or not with this enzyme (–), followed by protein separation by 8% SDS-PAGE gels (F) or 4-12% LDS-NuPAGE gels (G) in distinct running buffers, before being visualized by Western blotting with antibodies against V5 tag or USP19, respectively. The relevant images were cropped and also presented in the main Figure 3D.

#### Supplementary Figure S5

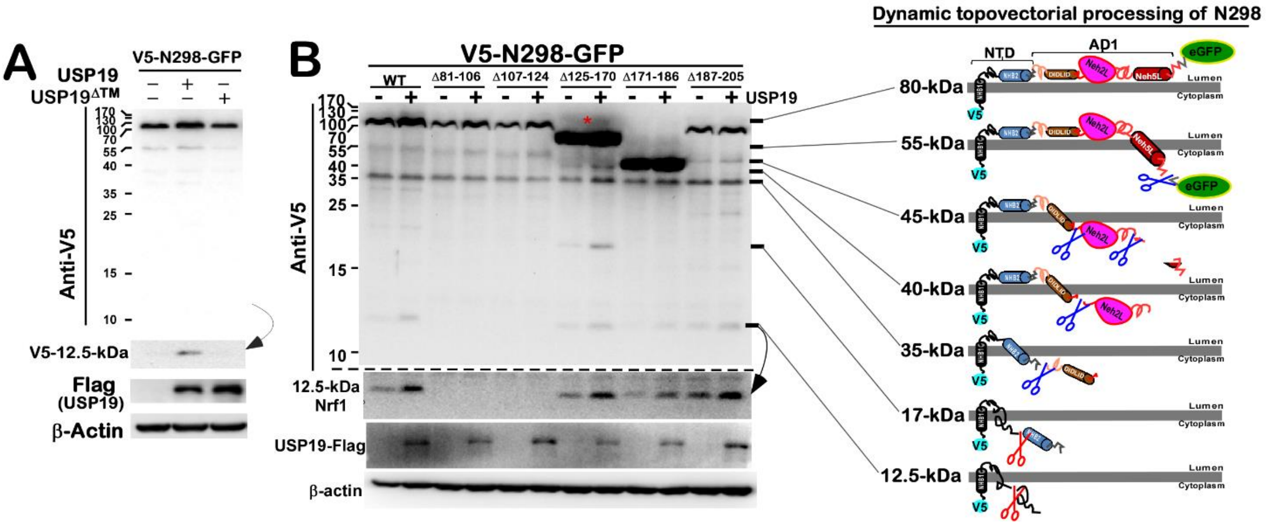


**Figure S5. Deletion of various N298 peptides leads to altered effects of USP19 on Nrf1.**

(A) A sandwiched chimeric protein, called V5-N298-GFP, were allowed for co-expression with USP19 (+), USP19$^{\Delta TM}$ (+), or neither (–), before immunoblotting with either V5 or USP19 antibodies. Among V5-N298-GFP, isoforms, its N-terminally V5-tagged 12.5-kDa images were enhanced and cropped from the upper same gels

(B) COS-1 cells, that had been allowed for ectopic expression of sandwiched chimeric protein V5-N298-GFP or one of its deletion mutants alone or plus USP19, were further evaluated by Western blotting with anti-V5 or -USP19 antibodies. The N-terminal 12.5-kDa images from processed V5-N298-GFP were enhanced and cropped from the upper same gels. In addition, the other relevant images were also presented in the main Figures 4B & 4C. Moreover, putative topological folding of V5-N298-GFP and its N-terminally-truncated polypeptides were, respectively, illustrated (in the *left panels*).

### Supplementary Figure S6

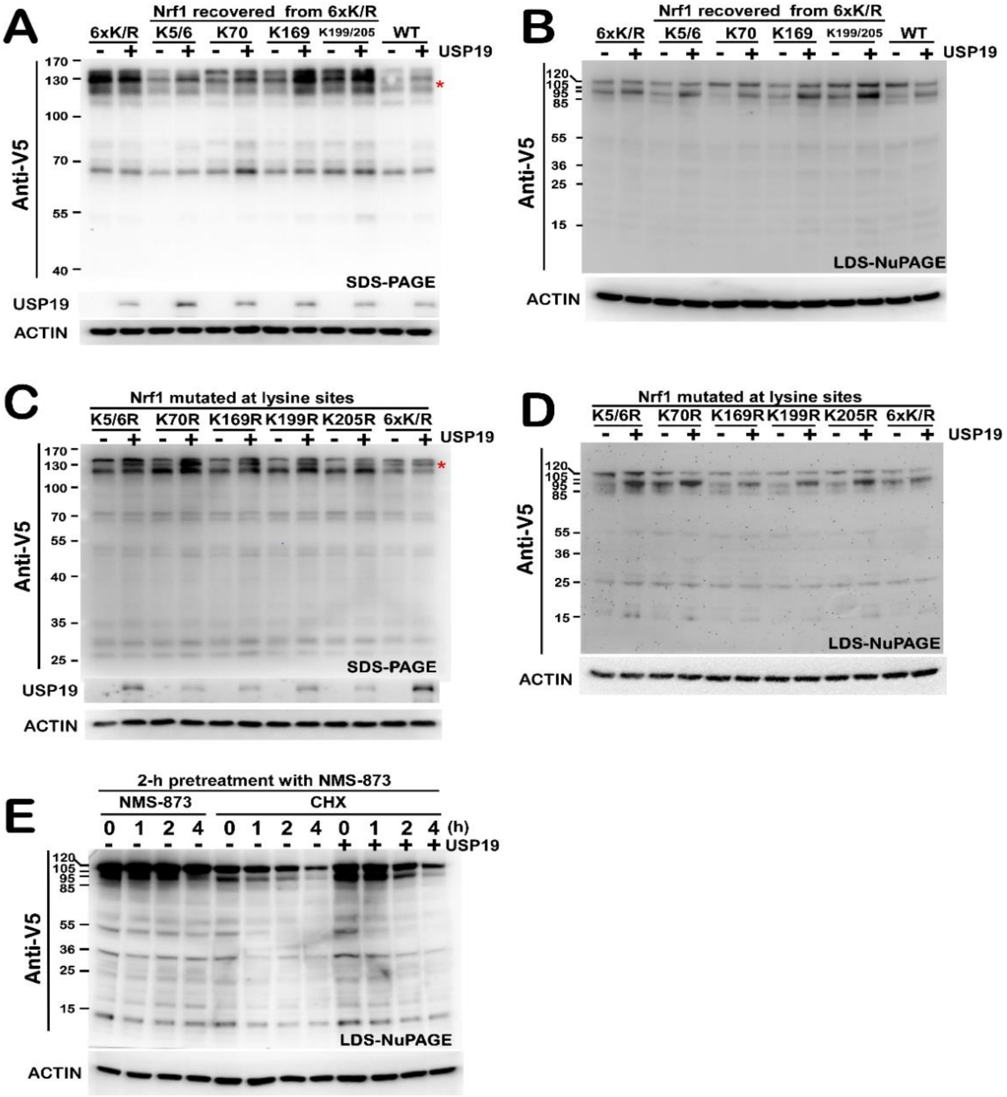


**Figure S6. Putative deubiquitination of Nrf1 by USP19 contributes to its stability and processing.**

(A,B) Total lysates of COS-1 cells, that had been transfected with expression constructs for wild-type Nrf1 (WT, 6×K), its mutant $Nrf1^{6\times K/R}$ (i.e., 6×K/R) or one of those recovered variants (i.e., K5/6, K70, K169 and K199/K205) from the $Nrf1^{6\times K/R}$ mutant, plus USP19 or not, were separated by 8% SDS-PAGE gels (A) or 4-12% LDS-NuPAGE gels (B) in distinct running buffers, and then visualized by immunoblotting with V5 or USP19 antibodies. The images cropped from the same gels are shown in Figure 4D.

(C, D) Western blotting of COS-1 cell lysates expressing one of distinct lysine-to-arginine mutants of Nrf1 (i.e. K5/6R, K70R, K169R, K199R, K205R and 6×K/R) alone or plus USP19 was carried out as described above. The images cropped from the same gels are shown in Figure 4E.

(E) COS-1 cells expressing Nrf1 alone or plus USP19 were pretreated for 2 h with NMS-873 (10 μmol/L) and then treated with CHX (50 μg/ml) for indicated lengths of time, before being harvested. Then total lysates were resolved by 4-12% LDS-NuPAGE gels and visualized by Western blotting as described above. The images cropped from the same gels are also shown in Figure 4F (*upper panel*).

### Supplementary Figure S7

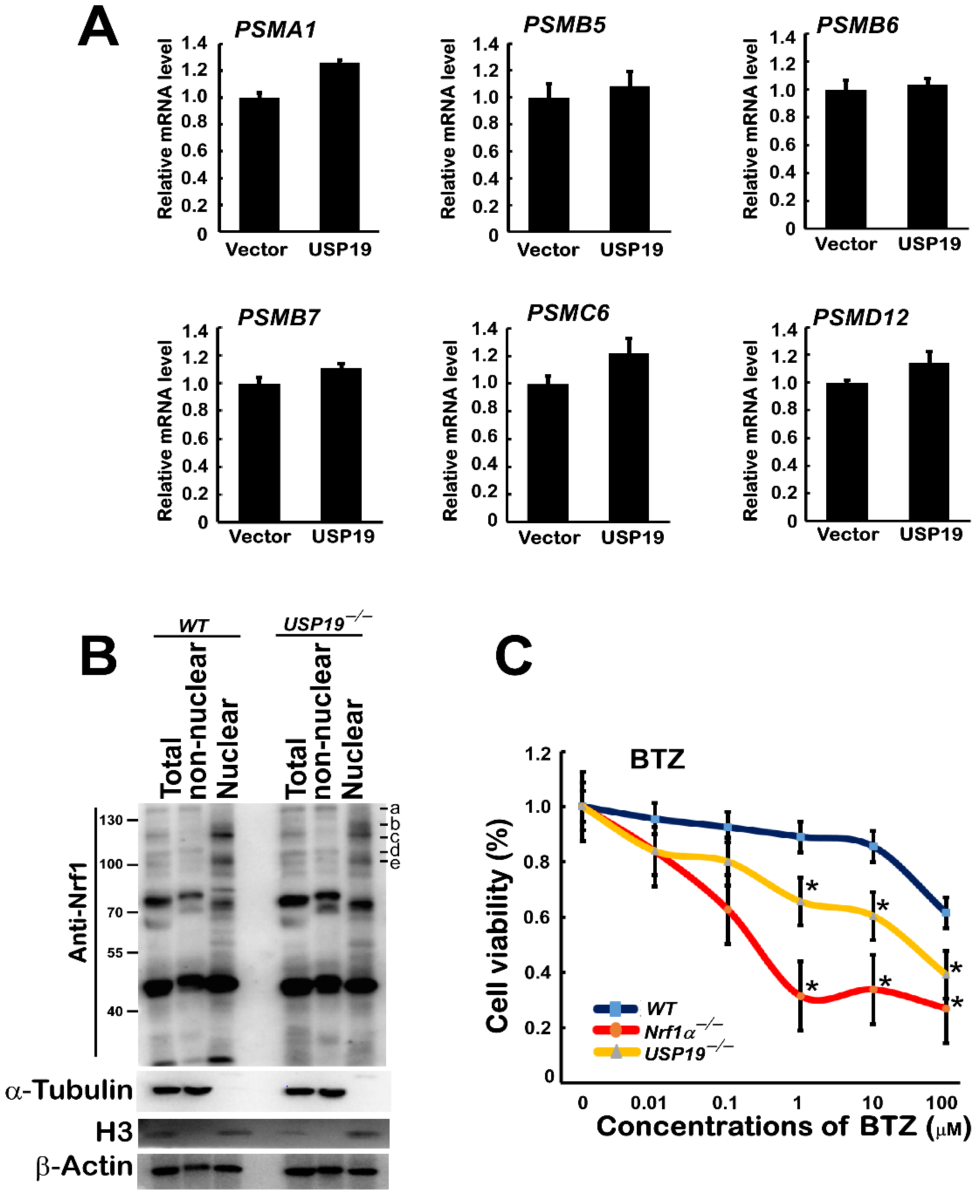


**Figure S7. USP19 potentiates the nuclear translocation of Nrf1 and its resistance to cytotoxity of BTZ.**

(A) HepG2 cells, that had been transfected with an expression construct for USP19 or empty vector, were examined by real-time qPCR analysis of some proteasomal subunits encoded by *PSMA1, PSMB5, PSMB6, PSMB7, PSMC6* and *PSMD12*. The resulting data are calculated as fold changes (mean ± SEM, $n = 3 \times 3$).

(B) Subcellular fractions of *WT* and *USP19*$^{-/-}$ cells were determined by immunoblotting with the indicated antibodies. The images cropped from the same gels are also shown in Figure 5C.

(C) Viability of *WT*, *Nrf1α*$^{-/-}$ and *USP19*$^{-/-}$ cell lines, that had treated for 24 h with different concentrations of bortezomib (BTZ). The resulting data are shown as fold changes (mean ± SEM, $n = 3 \times 3$) with significant decreases (* $p < 0.01$) as compared with the controls. The similar work had been done by Ze Zheng, as described by Zhu YP, *et al.* (in 2019 of the reference entitled 'Unification of Opposites between Two Antioxidant Transcription Factors Nrf1 and Nrf2 in Mediating Distinct Cellular Responses to the Endoplasmic Reticulum Stressor Tunicamycin', Antioxidants *9*, 4).

#### Supplementary Figure S8

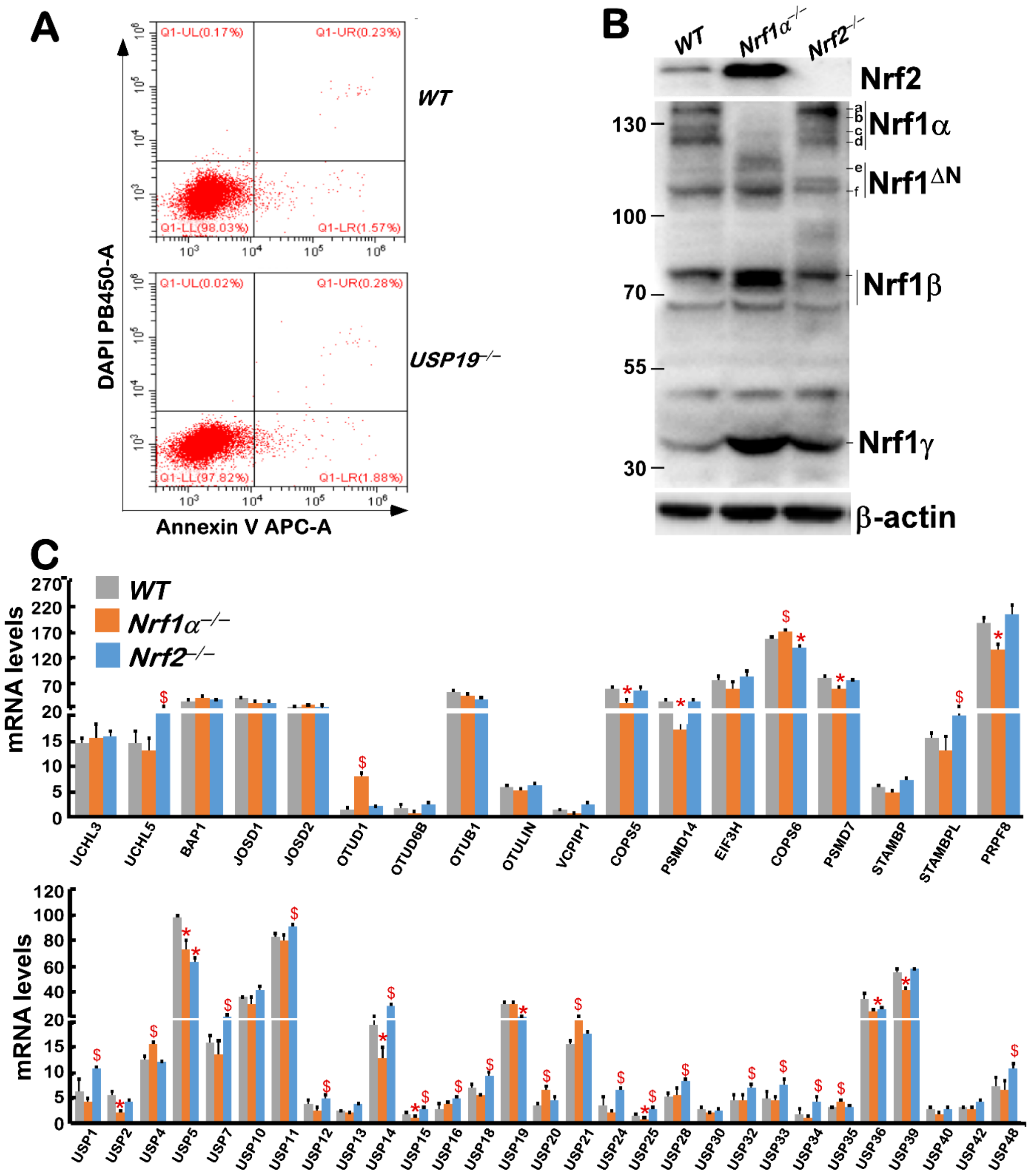


**Figure S8. Altered transcriptional expression of most deubiquitinases in *Nrf1α*$^{-/-}$ *and Nrf2*$^{-/-}$ cells.**

(A) Flow cytometry analysis of *WT* and *USP19*$^{-/-}$ cell apoptosis. UR and LR represent two fractions of early apoptotic cells and late apoptotic cells, whilst UL and LL denote another two fractions of necrotic cells and normal cells, respectively.

(B) Distinct abundances of Nrf1 and Nrf2 proteins in *WT*, *Nrf1α*$^{-/-}$ and *Nrf2*$^{-/-}$ cells were determined by Western blotting.

(C) Alterations in mRNA expression levels of most deubiquitinating enzymes in *Nrf1α*$^{-/-}$ and *Nrf2*$^{-/-}$ cells were determined by transcriptomic sequencing. The relevant data are calculated as fold FPKM values (mean ± SEM, $n$ = 3) with significant increases ($, $p < 0.01$) or significant decreases (* $p < 0.01$), when compared with the equivalents from *WT* control cells.

**Table1. A list of all key reagents and resources used in this work**

| Reagent or Resource | Source | Identifier |
|---|---|---|
| **antibodies** | | |
| Flag | Beyotime | AF519 |
| Nrf1 | Zhang's | N/A |
| Nrf2 | Abcam | Ab62352 |
| β-actin | ZSGB-BIO | TA-09 |
| Tubulin | Beyotime | AF0001 |
| USP19 | Abcam | Ab189518 |
| PSMB5 | Abclonal | A1975 |
| PSMB6 | Abclonal | A4053 |
| PSMB7 | Abclonal | A14771 |
| HA | Abcam | ab236632 |
| Histone 3 | Bioss | bs-0349R |
| ub | Cell Signaling Technology | 3933S |
| **Chemicals, Peptides, and Recombinant Proteins** | | |
| MG132 | Sigma Aldrich | M7449 |
| NMS-873 | selleck | S7285 |
| **Experimental Models: Cell Lines** | | |
| HepG2 | Cell bank of the Chinese Academy of Sciences | TCHu72 |
| COS-1 | Cell bank of the Chinese Academy of Sciences | GNO28 |
| $USP19^{-/-}$ | This paper | N/A |
| $Nrf1^{-/-}$ | Zhang's laboratory | N/A |
| $Nrf2^{-/-}$ | Zhang's laboratory | N/A |

| | | |
|---|---|---|
| **Recombinant DNA** | | |
| pARE-luc | Zhang's | N/A |
| pcDNA3.1 | Invitrogen | V79020 |
| p3XFLAG-CMV-14 | Sigma Aldrich | E7908 |
| pGL3-promoter | Promega | VQP0124 |
| pRL-TK | Promega | VQP0126Nrf |
| Nrf1-V5 | This paper | N/A |
| Nrf2-V5 | This paper | N/A |
| USP14-Flag | This paper | N/A |
| USP15-Flag | This paper | N/A |
| USP19-Flag | This paper | N/A |
| USP30-Flag | This paper | N/A |
| USP48-Flag | This paper | N/A |
| USP19-Luc | This paper | N/A |
| Mutants of USP19 | This paper | N/A |
| Mutants of Nrf1 | Zhang's laboratory | N/A |
| HA-Ub | BioVector | HA-ub pCMV |
| **Oligonucleotides for qPCR** | | |
| Nrf1 FW | Tsingke | GCTGGACACCATCCTGAATC |
| Nrf1 REV | Tsingke | CCTTCTGCTTCATCTGTCGC |
| Nrf2FW | Tsingke | TCAGCGACGGAAAGAGTATGA |
| Nrf2REV | Tsingke | CCACTGGTTTCTGACTGGATGT |
| PSMB5FW | Tsingke | GCTGTTGGCTCGGCAATGT |
| PSMB5 REV | Tsingke | GCCTCTCTTATCCCAGCCACAG |
| PSMB6 FW | Tsingke | TCAAGAAGGAGGGCAGGTGT |
| PSMB6 REV | Tsingke | GTAAAGTGGCAACGGCGAA |

| | | |
|---|---|---|
| PSMB7 FW | Tsingke | CTGTGTCGGTGTATGCTCCA |
| PSMB7 REV | Tsingke | TGCCAGTTTTCCGGACCTTT |
| ACTB FW | Tsingke | TGGCATCCACGAAACTACCTT |
| ACTB REV | Tsingke | CTTCTGCATCCTGTCGGCAAT |
| PSMA1 FW | Tsingke | ACAAATAAGCCCAACTTCAGG |
| PSMA1 REV | Tsingke | GCCAGCAAAATAATTTCTTCCC |
| PSMC6 FW | Tsingke | TACTGAAAATCCATGCAGGT |
| PSMC6 REV | Tsingke | GATCTGCTCCATTAAAGCC |
| PSMD12 FW | Tsingke | TTAAGTCCGCTCTATAGGGAT |
| PSMD12 REV | Tsingke | AGACTTACAAGTTAAAGCCACA |
| USP19 FW | Tsingke | AGCGAAGTATTGGACTCCCTC |
| USP19 REV | Tsingke | AGGAACACACGATGAAAACGA |
| **Oligonucleotides for construct** | | |
| USP14-Flag FW | Tsingke | GGAATTC ATGCCGCTCTACTCCGTTACTG |
| USP14-Flag REV | Tsingke | GGGGTACCTACTGTTCACTTTCCTCTTCC |
| USP15-Flag FW | Tsingke | CCCAAGCTTCAAGTTTGTACAAAAAAGTTG |
| USP15-Flag REV | Tsingke | TGCTCTAGAGTTAGTGTGCATACAGTTTTC |
| USP19-Flag FW | Tsingke | G GAATTC ATGTCTGGCGGGGCCAGTGC |
| USP19-Flag REV | Tsingke | GCTCTAGATCTCCAGCGACTCTGGGATAC |
| USP30-Flag FW | Tsingke | GGAATTCATGCTGAGCTCCCGGGCCGAGG |
| USP30-Flag REV | Tsingke | G GGG TAC CTATTCTTCAGACTTGCACTCCTG |
| USP48-Flag FW | Tsingke | GGAATTCATGGCCCCGCGGCTGCAGCTGGAG |
| USP48-Flag REV | Tsingke | GGGGTACCTAATGTCCAAGAAGACCAGTACC |
| Nrf1-V5 FW | Tsingke | GGGGTACCATGCTTTCTCTGAAGAAATACTTAAC |
| Nrf1-V5 REV | Tsingke | GCTCTAGACTCTTTCTCCGGTCCTTTGGCTTCCTC |
| Nrf2-V5 FW | Tsingke | GGGGTACCATGATGGACTTGGAGCTGCCG |

| | | |
|---|---|---|
| Nrf2-V5 REV | Tsingke | GCTCTAGACTGTTTTTCTTAACATCTGGCTTC |
| USP19-Luc FW | Tsingke | GGGGTACCGGCCTCCTCTTGACTACTGTGCTCATC |
| USP19-Luc REV | Tsingke | GCAGCGCAAGCTTACTGACCACAGATCA |
| Software and Algorithms | | |
| Canvas X | Canvas GFX, Inc. | https://www.canvasgfx.com/ |
| FlowJo 7.6.5 | FlowJo | https://www.flowjo.com/ |
| Primer Premier 5 | PREMIER Biosoft International | https://www.PremierBiosoft.com/ |
| CFX Manager 3.1 | Bio-Rad | https://bio-rad-cfx-manager.com/ |
| Quantity One | Bio-Rad | https://bio-rad-cfx-manager.com/ |
